# Distinct Colitis-Associated Macrophages Drive NOD2-Dependent Bacterial Sensing and Gut Homeostasis

**DOI:** 10.1101/2025.01.21.634180

**Authors:** Gajanan D. Katkar, Mahitha Shree Anandachar, Stella-Rita Ibeawuchi, Ella McLaren, Megan Estanol, Kennith Carpio-Perkins, Shu-Ting Hsu, Celia R. Espinoza, Jane Coates, Yashaswat S. Malhotra, Madhubanti Mullick, Vanessa Castillo, Daniella T. Vo, Saptarshi Sinha, Pradipta Ghosh

## Abstract

Single-cell studies have revealed that intestinal macrophages maintain gut homeostasis through the balanced actions of reactive (inflammatory) and tolerant (non-inflammatory) subpopulations. How such balance is impaired in inflammatory bowel diseases (IBD), including Crohn’s disease (CD) and ulcerative colitis (UC), remains unresolved. Here, we define colon-specific macrophage states and reveal the critical role of *<u>n</u>*on-*i*nflammatory *<u>c</u>*olon-*<u>a</u>*ssociated *<u>m</u>*acrophages (niColAMs) in IBD recovery. Through trans-scale analyses—integrating computational transcriptomics, proteomics, and *in vivo* interventional studies—we identified GIV (*CCDC88A*) as a key regulator of niColAMs. GIV emerged as the top-ranked gene in niColAMs that physically and functionally interacts with NOD2, an innate immune sensor implicated in CD and UC. Myeloid-specific GIV depletion exacerbates infectious colitis, prolongs disease, and abolishes the protective effects of the NOD2 ligand, muramyl dipeptide, in colitis and sepsis models. Mechanistically, GIV’s C-terminus binds the terminal leucine-rich repeat (LRR#10) of NOD2 and is required for NOD2 to dampen inflammation and clear microbes. The CD-associated *1007fs* NOD2-variant, which lacks LRR#10, cannot bind GIV—providing critical insights into how this clinically relevant variant impairs microbial sensing and clearance. These findings illuminate a critical GIV-NOD2 axis essential for gut homeostasis and highlight its disruption as a driver of dysbiosis and inflammation in IBD.

## INTRODUCTION

Intestinal macrophages are critical for gut development, immunity and repair (1). Single-cell studies have revealed that gut homeostasis relies on the dynamic interplay between two antagonistic macrophage subpopulations: inflammatory “accelerators” and non-inflammatory “brakes” (2, 3). An imbalance in these subpopulations can lead to uncontrolled gut inflammation, as observed in inflammatory bowel diseases (IBD) such as Crohn’s disease (CD) and ulcerative colitis (UC) (4, 5). However, precisely defining these subpopulations and understanding their roles in health and disease, and the molecular mechanisms that control the same, remain significant challenges (6).

Recent advances in artificial intelligence (AI) and machine learning-guided transcriptomics have addressed this challenge by enabling the analysis of diverse macrophage states across both bulk and single-cell datasets (3, 7–10). Among these approaches, Boolean implication networks have emerged as a robust method with a decade-long track record (7, 8, 11, 12) for identifying universally conserved gene expression patterns (or “invariants”). These patterns remain consistent despite the variability introduced by tissue heterogeneity, circadian rhythms, metabolic states, species diversity, perturbations, stimuli, and disease conditions (7, 8, 11, 12). Using this network approach on a dataset of pooled isolated monocytes/macrophages representing greatest possible diversity, we recently defined ‘*<u>S</u>*ignature of *<u>Ma</u>*crophage *<u>R</u>*eactivity and *<u>T</u>*olerance’ (SMaRT; see **Figure 1A**, Supplemental information 1) as a conserved 338-gene signature representing macrophage continuum states, across the physiologic and pathologic spectra of “reactivity" and "tolerance" (3). We showed that while the conventional M1/M2 classification fails to capture the diversity, plasticity, and continuum of macrophage states in tissue during homeostasis and disease, the SMaRT model-derived definitions remain robust and consistently outperform other emerging classification schemes across contexts (3).

**Figure 1.**
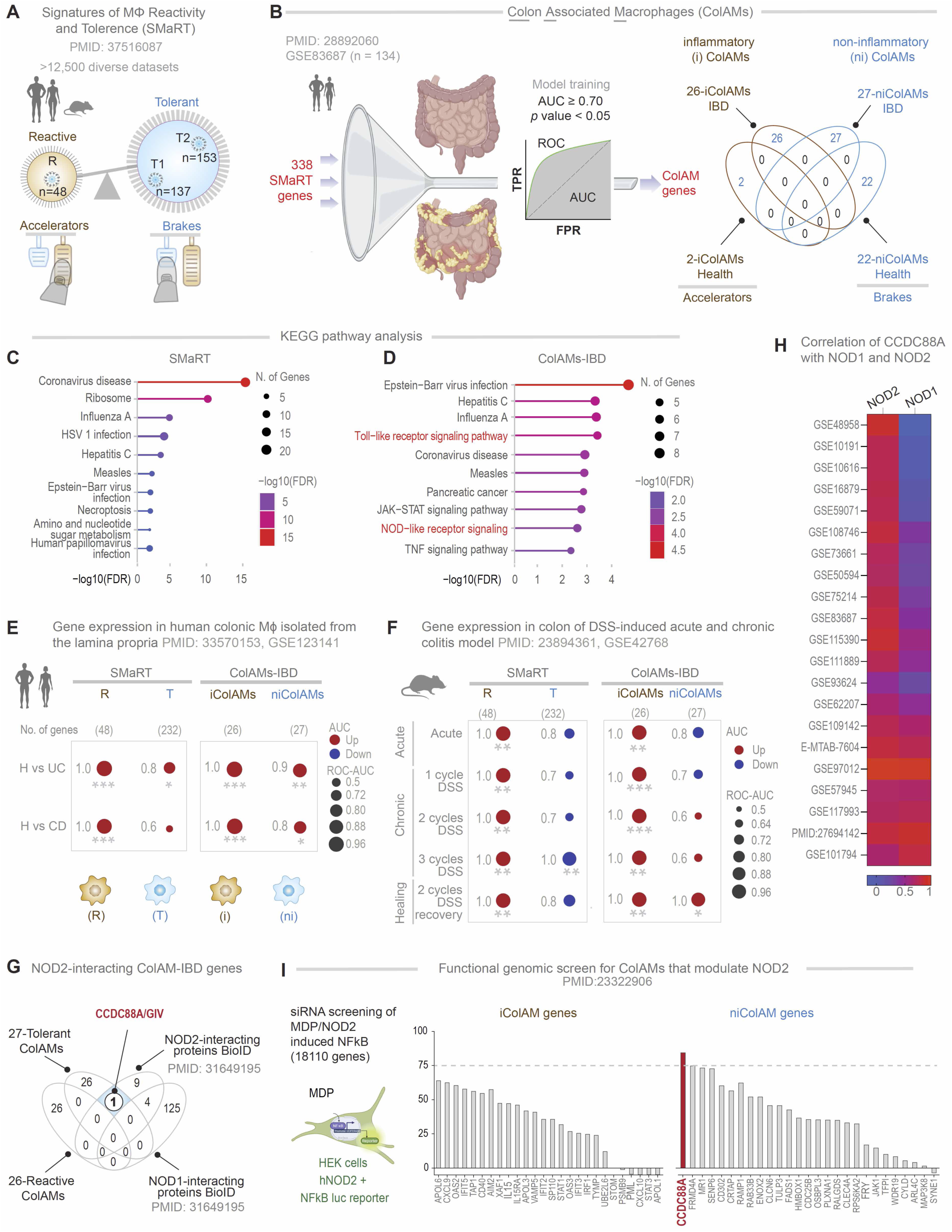
Identification of *CCDC88A* as a putative modulator of NOD2 in non-inflammatory ColAMs in IBD. **A**. Key steps of the previously published workflow (3) used to develop the computational model of macrophage continuum states—*<u>S</u>*ignatures of *<u>Ma</u>*crophage *<u>R</u>*eactivity and *<u>T</u>*olerance (SMaRT)—are shown. SMaRT identifies invariant gene clusters representing reactive (R) and tolerant (T1, T2) states across over 12,500 diverse transcriptomic datasets. The schematic below illustrates their opposing roles: reactive macrophages act as “accelerators,” while tolerant states serve as “brakes,” working antagonistically to fine-tune inflammatory responses to perceived threats. **B**. *<u>Left</u>*: Key steps used here to refine SMaRT in the context of the gut mucosa and derive colon-associated macrophage signatures (ColAMs) using a dataset (<u>GSE83687</u>) (24) comprised of both healthy and IBD samples. SMaRT genes were trained to derive a subset of ColAMs that could classify healthy vs IBD sample, achieving an area under the curve ≥ 0.7. *<u>Right</u>*: Inflammatory and non-inflammatory ColAMs identified here, in health (top) and IBD (bottom). **C-D.** KEGG pathway enrichment analysis on the SMaRT (**C**) and the ColAM-IBD (**D**) genes. **E-F.** Bubble plot displays ROC-AUC values (circle radii proportional to ROC-AUC) illustrating gene regulation directions (Up-regulated in red, Down-regulated in blue) for classifying healthy vs. CD and healthy vs. UC in human colonic lamina propria (<u>GSE123141</u>) (**E**) and DSS-induced acute, chronic and healing phase of murine colitis models (**F**). This classification is based on macrophage gene signatures of reactivity ‘R’ and tolerance ‘T’, identified in the SMaRT model and the i/niColAMs, used independently. Welch’s two-sample unpaired t-test is applied to the composite gene signature score (z-score of normalized transcripts per million [tpm] count) to compute p-values (* *p* ≤ 0.05; ** *p* ≤ 0.01; *** *p* ≤ 0.001). **G.** Venn diagram shows the number of genes/proteins, identified by independent studies, to interact with NOD1/2. One gene/protein (*CCDC88A*/Girdin; white circle) interacts with NOD2 but not NOD1, is associated with tolerant ColAMs. **H**. Heatmap of the correlation coefficient of normalized gene expression of *CCDC88A* with NOD2 (left) and NOD1 (right) in various independent transcriptomic datasets of healthy and IBD tissues. **I**. The bar graph shows MDP/NOD2-induced NFκB activation observed during a functional genomic (siRNA-based) screen. The impact of depletion of iColAM genes and niColAM genes are presented. Interrupted line marks 75% enhancement of NFkB activity compared to MDP-stimulated control samples.

Here we sought to refine the SMaRT model in the context of IBD. We hypothesized that these definitions would yield robust classification and functional insights into the colitic environment. First, we formally define two macrophage subpopulations in the colon: inflammatory (iColAMs) and non-inflammatory (niColAMs) ---both in health and IBD. We find that tolerant niColAMs are essential for dampening inflammation and resolving infections, making them critical for recovery from IBD. We subsequently identify a previously unappreciated yet consequential physical and functional coupling in IBD-associated niColAMs between the innate immune sensor nucleotide-binding oligomerization domain-containing protein 2 (NOD2) and GIV (Gα-interacting vesicle-associated protein, also known as Girdin). NOD2, also known as NLRC2, belongs to the nucleotide-binding domain and leucine-rich repeat (NLR) family and functions as an intracellular pattern recognition receptor (PRR) for muramyl dipeptide (MDP) derived from pathogens. NOD2 coordinates bacterial clearance and confers immunity (4), all while mounting a controlled inflammatory program that involves the dampening of NF-κB (13) activity that is TLR2/4-dependent (14–18). GIV, on the other hand, is a multimodular signal transducer and the prototypical member of the non-receptor *<u>G</u>*uanine nucleotide *<u>E</u>*xchange *<u>M</u>*odulator (GEM (19)) family of proteins. Unlike the canonical GPCR/G protein pathway, in which G proteins engage exclusively with ligand-activated GPCRs, GEMs like GIV bind and modulate G protein activity downstream of a myriad of cell-surface receptors (20–22). Of relevance here, GIV is a ubiquitously expressed molecule that is highly expressed in immune cells such as macrophages and serves as a “brake” for the cell surface PRR, TLR4 and modulates macrophage inflammatory responses to LPS (22) and gut barrier integrity during aging (23), cancer (23) and in IBD (7), and it’s gene (*CCDC88A*) has emerged as a key determinant of macrophage polarization in the SMaRT model (3). We demonstrate that GIV interacts dynamically with NOD2 to facilitate microbial sensing and clearance while also suppressing inflammation. This protective mechanism is disrupted in the most clinically significant IBD-associated NOD2 risk variant, highlighting its relevance to disease pathology. These insights shed new light on the molecular pathways underlying gut homeostasis and the progression of IBD, offering potential therapeutic avenues for restoring balance in macrophage subpopulations.

## RESULTS

### Identification of distinct subpopulations of colon-associated macrophages

To contextualize the SMaRT model (**Figure 1A**, Supplemental information 1) within the human gut, and specifically, in IBD, we refined it using the largest, high-quality, full-thickness colon tissue transcriptomic dataset available for IBD (<u>GSE83687)</u> (24)--the only dataset of its kind. Because the original model was built using purified macrophages and monocytes from diverse tissues, we assumed that refinement using bulk RNA-seq data would preserve a subset of macrophage-specific genes from the SMaRT model that are most relevant to IBD. Briefly, we employed a machine learning-based classifier on 338 SMaRT signature genes (3) (Supplemental information 1) to identify the classification accuracy of each of the SMaRT signature genes on healthy vs IBD-affected colon tissues (**Figure 1B**). This allowed us to formally define ColAMs in health as those expressing a core set of 24 genes [2, that are expressed highly in reactive inflammatory (i) ColAMs; 22 that are expressed highly in tolerant non-inflammatory (ni)ColAMs; **Figure 1B**] and in IBD, as those expressing a distinct set of 53 genes (26, that are expressed highly in reactive iColAMs; 27, that are expressed highly in tolerant niColAMs; **Figure 1B**). It is noteworthy that the brakes and accelerators in health are distinct from IBD (**Figure 1B***, see* Supplemental information 2). The ColAM genes had AUC values greater than 0.70 (**Figure 1B**, Supplemental information 2) in discriminating between healthy and IBD-affected colon tissues (including UC and CD, see Supplemental information 2). KEGG pathway enrichment analysis revealed that model refinement led to enrichment of colitis-relevant pathways— including Toll-like receptor, NOD2, and TNF signaling (compare **Figure 1, C** and **D;** see Supplemental information 2 for gene lists).

When we tested their ability to distinguish healthy from colitis samples, the 53-gene ColAM signature (used independently as 26-gene iColAMs and 27-gene niColAMs) performed consistently better than the original SMaRT model (3), in both human (H vs. UC/CD; **Figure 1E**) and murine (H vs. dextran sodium sulfate (DSS), a chemical colitogen; **Figure 1F**) datasets. Leveraging a high-quality murine dataset of DSS-induced acute and chronic colitis (25), we found that i/niColAMs may be indued in temporally distinct patterns; iColAMs were induced acutely and persisted throughout the various DSS models, whereas niColAMs were induced exclusively in chronic model in which injury was repetitive in the form of 2 cycles of DSS followed by 3 weeks of recovery/washout, which is believed to better recapitulate the relapsing-remitting nature of IBD, (**Figure 1F**).

### NOD2 may functionally couple with CCDC88A in colitis-associated non-inflammatory ColAMs

NOD2, located on chromosome 16, remains the most replicated genetic association in IBD, with a mean allelic odds ratio of 3.1 across studies (26, 27), and a well-established, though mechanistically debated, role in IBD pathogenesis (4, 28–32). Persistent controversy surrounds how NOD2 functions and how its variants drive colitis in both UC and CD (32–36). Given the enrichment of NOD signaling in IBD-associated colonic macrophages (**Figure 1D**) we investigated NOD-centric cellular processes in i/niColAMs.

Overlaying i/niColAM gene clusters with a published NOD1/2 interactome, as determined by BioID Proximity-Dependent Biotin Identification; (37), we identified a single candidate interactor: GIV, encoded by *CCDC88A* (**Figure 1G**). Notably, *CCDC88A* is part of the niColAM gene signature, which emerges during the recovery phase of DSS-induced colitis (**Figure 1F**). Its expression correlates with NOD2—but not NOD1—across 21 independent cohorts (**Figure 1H**) and is elevated in intestinal macrophages from UC and CD patients compared to healthy controls (**Figure S1A**).

We next leveraged a genome-wide siRNA screen in HEK293T cells (38) that assessed MDP-induced hyperactivation of NFκB. While NOD2 variants are known to impair bacterial clearance and disrupt NFκB activation (13, 29, 38, 39), paradoxically, the gut mucosa of IBD patients often shows heightened NFκB activity (40–44). Loss-of-function NOD2 variants, such as the CD-associated *1007fs* (45), are also known to impact the severity of disease course in UC (46). Based on these observations, NOD2 is believed to restrict activation of the NFκB pathway by TLR2/4 (14–18) and its dysfunction causes runaway inflammation, thereby increasing the risk of colitis. Consistent with these observations, the functional-genomic screen revealed that among all i/niColAM-genes, depletion of *CCDC88A* within the niColAM cluster (genes presumed to be critical for reducing inflammation) emerged as the most consequential perturbation that increases NFkB activity (**Figure 1I**).

These findings suggest that CCDC88A may functionally couple with NOD2 to restrain inflammation in colonic macrophages, providing a strong rationale to investigate the protective niColAM state during colitis recovery.

### GIV is required for MDP/NOD2-mediated bacterial clearance and controlled inflammation

Given its recently identified role in modulating macrophage responses (22), we asked if GIV may be a functional modulator of the cytosolic sensor, NOD2. To study the role of GIV in MDP/NOD2-induced inflammatory responses in macrophage *in vitro*, we used 4 cell-based models: (i) GIV-depleted (shGIV) RAW 264.7 murine macrophage cells; this previously-validated cell model displays ∼85-90% depletion of GIV protein by immunoblotting (22) (**Figure 2A**)]; (ii) THP1 NFkB-SEAP reporter human macrophage lines depleted or not of >∼90% GIV protein (by CRISPR; **Figure 2C**). (iii) thioglycolate-induced murine peritoneal macrophages (TGPMs) isolated from myeloid-specific conditional GIV knockout (GIV-KO) mouse, generated previously (22) by crossing Girdin floxed mice to *LysM*cre mice, and confirmed to have ∼85-90% depletion of GIV protein; and (iv) THP1 human macrophage lines depleted or not of >∼90% GIV protein (by CRISPR; Figure 2F).

**Figure 2.**
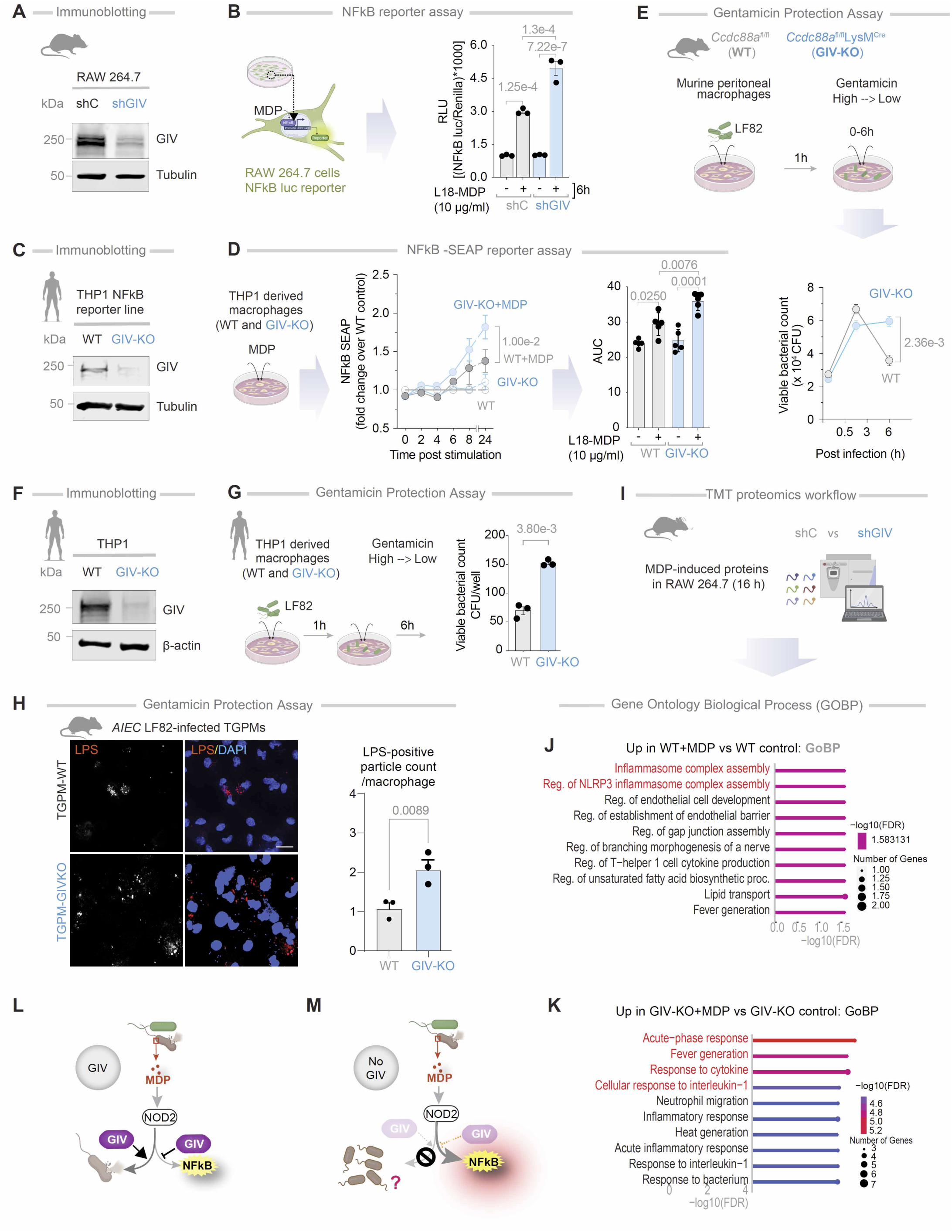
GIV is required for dampening MDP-stimulated inflammation and promoting bacterial clearance in macrophage. **A.** Immunoblot of control (shC) or GIV-depleted (shGIV) RAW 264.7 cells. **B.** Workflow of the NFκB reporter assay in RAW 264.7 cells (*left*). Bar graphs display the fold change in NFκB activity (*right*). **C.** Immunoblot of control (WT) and GIV-knockout (GIV-KO) THP1-NFkB SEAP reporter cell line-derived macrophages. **D.** Workflow of the NFκB reporter assay in CRISPR-depleted human THP-1 cells expressing an NFκB activity-tracking reporter (*left*). Line graphs display the fold change in NFκB activity over WT control (*right*). **E.** Schematic (top) displays the workflow for bacterial clearance assay. Line graphs (below) show the viable bacterial counts in the peritoneal macrophages. **F.** Immunoblot of control (WT) or GIV-knockout (GIV-KO) THP1 monocyte derived macrophage. **G.** Workflow of the gentamicin protection assay in THP1 cells. Bar graphs show the viable bacterial counts in the THP1 monocyte derived macrophage. **H.** Immunofluorescence images display representative fields of TGPMs challenged with live *AIEC*-LF82 (MOI 1:30) for 1 h. Scale bar = 20 µM. Bar graphs display quantification of intracellular *AIEC*-LF82; n = 4-6. **I-K**: Workflow for multiplexed proteomics analyses (I). Bar graphs showing biological process as determined by gene ontology biological process analysis (Red, pathways cited in text). **L-M**. Schematic summarizing findings in cells with GIV (L) and without GIV (M). *Statistics*: All results are displayed as mean ± SEM (n = 3 biological replicates). Significance was tested using two-way/one-way ANOVA followed by Tukey’s test for multiple comparisons. *p*-value ≤ 0.05 is considered as significant.

In GIV-depleted RAW macrophages, MDP/NOD2-induced NFκB activity was significantly elevated (**Figure 2B**), as determined by luciferase reporter assays. Dynamic NFκB reporter assays in THP1 reporter cells further confirmed the findings, adding robustness to the results (**Figure 2D**). These findings were corroborated in HeLa cells (**Figure S1, B and C**), a cell line commonly used to study NOD2-dependent processes in plasmid transfection settings (47, 48). Briefly, compared to control cells, GIV-depleted HeLa cells (by CRISPR; **Figure S1B**) showed significantly higher MDP/NOD2-induced NFkB activity, confirming the role of GIV in dampening NFkB activity. Consistent with its role in dampening inflammation, GIV depletion in TGPMs led to a ∼3-4-fold increase in proinflammatory cytokines [interleukin (IL)1β, IL6, and tumor necrosis factor (TNF)α], as measured by ELISA (**Figure S1D**). Hyper-induction of proinflammatory cytokines was accompanied by a concomitant suppression of the anti-inflammatory cytokine, IL10 (**Figure S1D**). These cytokine profiles were consistent with gene expression patterns assessed via qPCR (**Figure S1E**).

When TGPMs were infected with *<u>a</u>*dherent-*i*nvasive *<u>E</u>scherichia <u>c</u>oli* strain-LF82 (*AIEC*-LF82), isolated from CD patients (49), GIV-KO TGPMs exhibited delayed bacterial clearance compared to WT controls (**Figure 2E**). Similarly, GIV-KO THP1 cells reproduced these findings (**Figure 2G**). Immunofluorescence imaging further confirmed that GIV-KO TGPMs retained significantly higher numbers of pathogenic *AIEC*-LF82 bacteria (**Figure 2H**).

Tandem mass tag (TMT)-based quantitative proteomics of GIV-depleted RAW macrophages revealed distinct proteomic differences after 16 hours of MDP stimulation (**Figure 2I**). While control cells activated robust NOD2-dependent signalling and inflammasome assembly (**Figure 2J**), GIV-depleted cells showed an acute-phase response and heightened expression of proinflammatory cytokines (**Figure 2K**).

Together, these results identify GIV as a critical mediator of MDP/NOD2 signaling. GIV is essential for maintaining a balanced pro- and anti-inflammatory cytokine response and promoting effective bacterial clearance. In its absence, macrophages exhibit exaggerated NFκB-driven inflammation but fail to clear bacteria efficiently (**Figure 2, L-M**), suggesting that GIV’s role in microbial clearance may be independent of its modulation of NFκB signaling.

### GIV is required for phagolysosomal fusion

To understand why GIV-deficient macrophages retain higher intracellular bacterial loads despite heightened NFκB activity (**Figure 2M**), we next investigated whether GIV plays a direct, NFκB-independent role in bacterial clearance. Because NOD2-dependent response to degraded bacteria requires the phagosomal membrane potential and the activity of lysosomal proteases (50), we hypothesized that GIV may facilitate phagolysosomal fusion.

We used two complementary approaches to test this. First, we challenged TGPMs in vitro with *AIEC*-LF82 and assessed the spatial proximity of internalized bacteria to LAMP1-positive lysosomes by confocal immunofluorescence microscopy (**Figure 3A**). In control (WT) cells, *AIEC*-LF82 bacteria were frequently found near LAMP1-positive structures (**Figure 3A**), suggesting efficient delivery of phagosomes to lysosomes. By contrast, GIV-deficient macrophages showed a marked reduction in bacteria-lysosome proximity (**Figure 3A**), suggesting disrupted lysosomal targeting.

**Figure 3.**
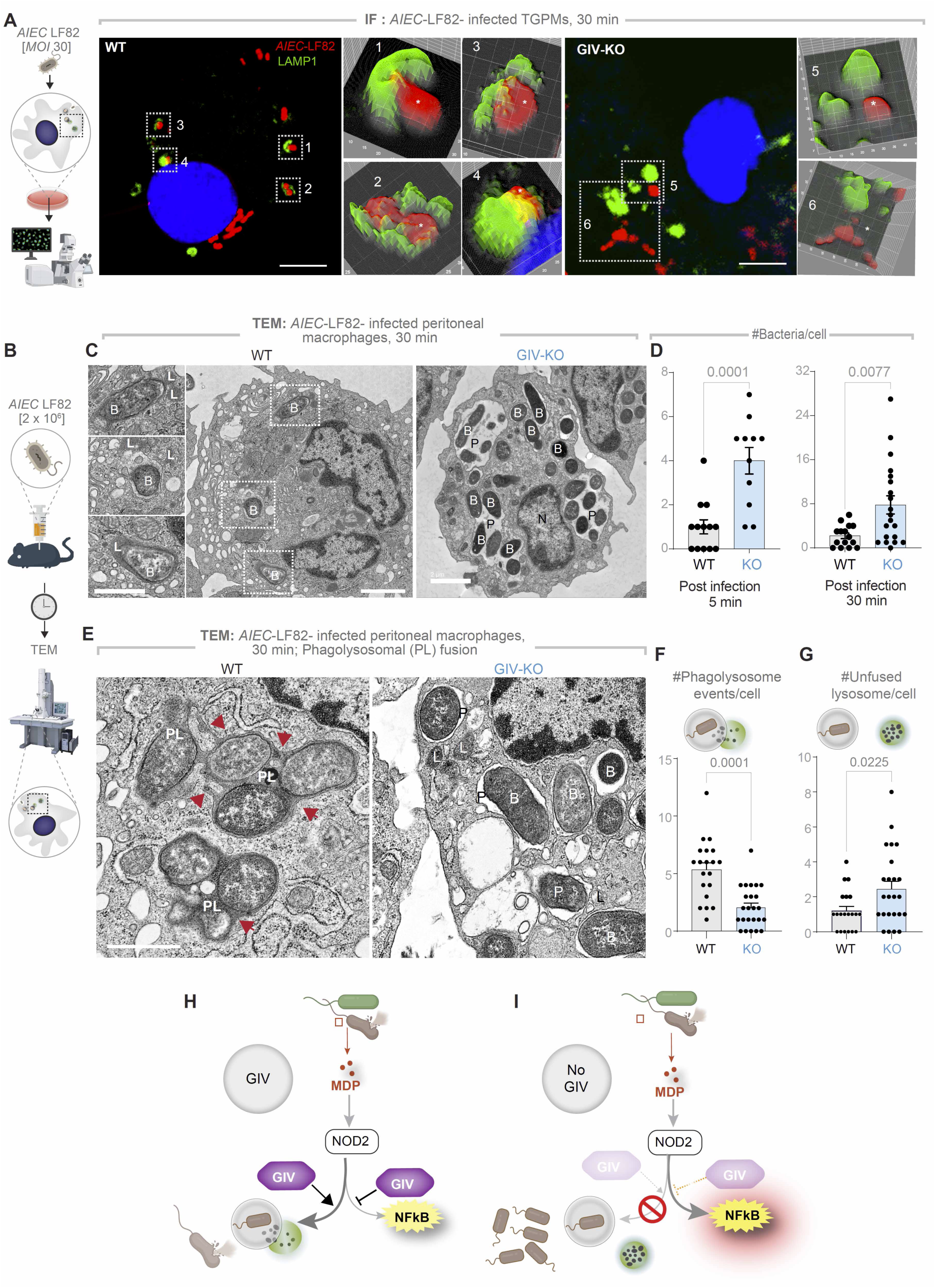
GIV is required for phagolysosomal fusion and bacterial clearance. **A.** Workflow for immunofluorescence studies of *AIEC*-LF82-challenged TGPMs (left) and representative images (right) showing the proximity of *AIEC*-LF82 (red) to LAMP1-positive lysosomes (green). Insets show magnified, 3D-rendered versions of boxed regions, created using ImageJ. Scale bars = 5 µm **B.** Workflow (B) for TEM studies on infected peritoneal macrophages in C-G. **C.** Representative TEM images showing bacterial abundance. Scale bar = 2 µm. **D.** Bar graphs quantifying the number of bacteria per cell at 5- and 30-min post-infection. **E.** High-magnification TEM images highlighting phagolysosomal (PL) fusion events (arrowheads). **F-G**. Bar graphs display the number of events per cell (F) and number of unfused lysosomes per cell (G); n = 2 repeats. N=nucleus, P=phagosome, L=lysosome, PL=phagolysosome, B= bacteria AIEC-LF82. **H-I.** Summary of findings in cells with (H) or without (I) GIV. *<u>Statistics</u>*: All TEM quantifications are based on ∼20-30 fields; n = 2 independent biological repeats. Results are presented as mean ± SEM. Significance was determined using Student’s t-test; p-value ≤ 0.05 was considered significant.

Second, we used quantitative transmission electron microscopy (TEM) to visualize phagolysosomal fusion events and quantify bacterial burden over time (**Figure 3B**). GIV-deficient macrophages harbored visibly higher numbers of intracellular *AIEC*-LF82 compared to WT controls (**Figure 3C**), both at 5- and 30-minutes post-infection (**Figure 3D**), confirming impaired bacterial clearance. TEM imaging also revealed stark ultrastructural differences: while WT cells exhibited numerous phagolysosomal (PL) fusion events (**Figure 3E**, arrowheads). GIV-deficient macrophages showed markedly fewer fusion events (**Figure 3F**), and retained a higher number of unfused lysosomes (**Figure 3G**), suggesting a defect in phagosome maturation and lysosome engagement, but not lysosome biogenesis.

These findings define a mechanistically distinct, NFκB-independent role for GIV in promoting phagolysosomal fusion. In its absence, bacterial clearance fails despite heightened inflammatory signaling, underscoring GIV’s dual function: restraining inflammation via NFκB modulation and promoting pathogen elimination through lysosomal trafficking (**Figure 3, H-I**).

### GIV-KO mice develop dysbiosis and exacerbated and protracted *Citrobacter*-induced colitis

To investigate the role of GIV in vivo, we employed a myeloid-specific GIV-KO (*Ccdc88a*^fl/fl^/*LysM*^Cre^) model (see *Methods*) (22). We found that these mice spontaneously develop dysbiosis by ∼8-12 wk (**Figure 4, A and B**; **Figure S2**). Notably, the strain *Rhizobiales —*uniquely associated with CD patients and absent in healthy controls (P = 0.037) (51) *—*detected in 100% of GIV-KO mice (5/5) but was undetectable in control littermates (**Figure 4, A and B**; **Figure S2**).

**Figure 4.**
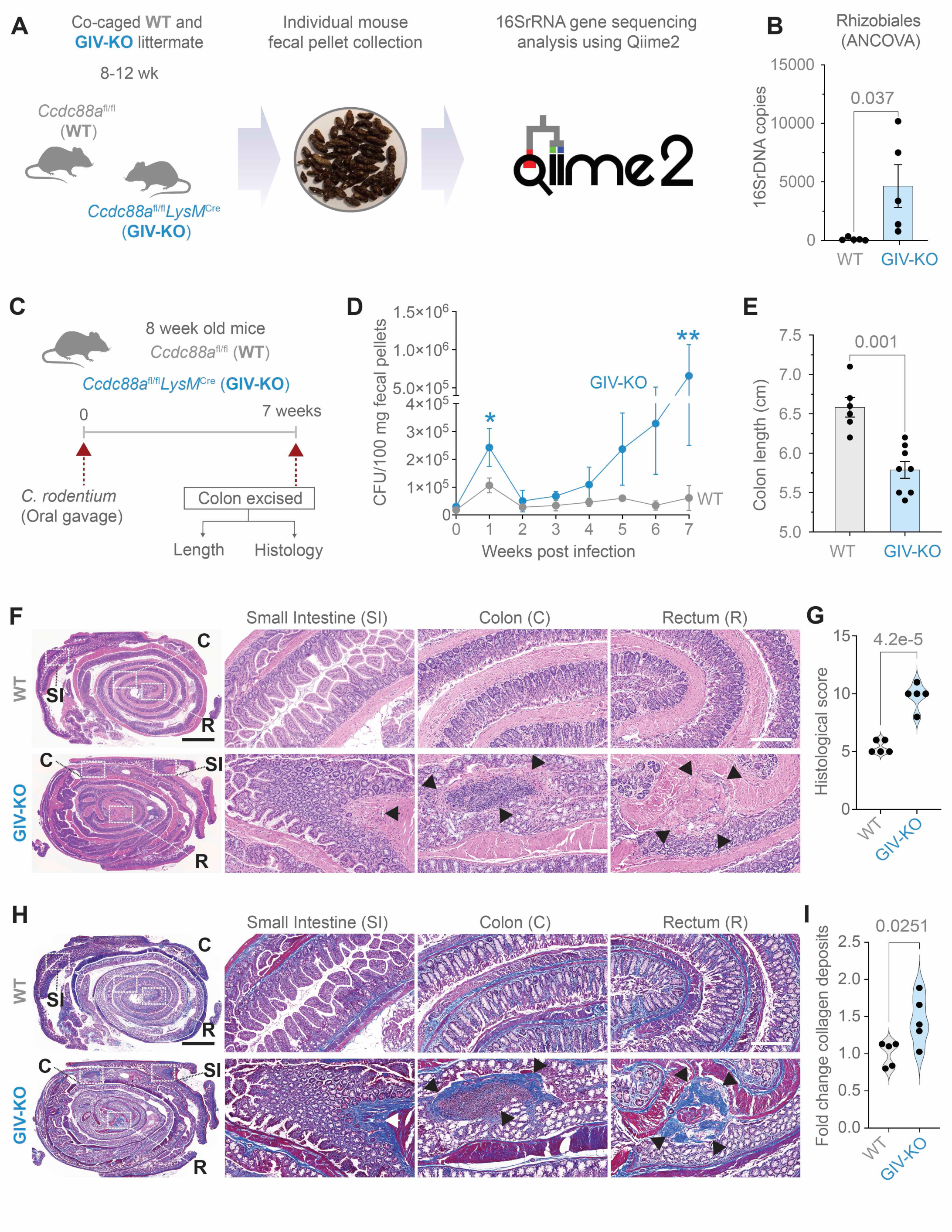
A mouse model of dysbiosis, impaired microbial clearance, patchy chronic transmural ileocolitis and fibrosis. **A-B**. Schematic (A) and bar graph (B) display the process and outcome of a 16S fecal microbiome analysis at baseline in 10 wk-old myeloid-specific (*LysM*Cre) GIV-KO mice and their control littermates (WT). *p* value was estimated by unpaired t-test; n = 5 mice in each group. **C-I**. Panels describing the experimental design (**C**) and findings (**D-I**) in an infectious colitis model of GIV-KO and control littermates induced using *Citrobacter rodentium* (initially termed *Citrobacter freundii* biotype 4280 (115); strain name DBS100; 5 x 10^8^ CFU/200ul/mouse. GIV-KO, n=8; WT, n=6. Findings are representative of two independent repeats. Line graphs (**D**) display the bacterial burden in fecal pellets over a 7 wk period after the initial oral gavage. P values were determined by unpaired t-test. *, < 0.05; **, 0.01. Bar graph (**E**) displays the differences in colon length. *p* values were determined by unpaired t-test. H&E (**F**) or trichrome (**H**)-stained images representative of Swiss rolls of the entire intestinal tract are shown. Scale bar = 2.5 mm. Magnified fields of the rectum (R), colon (C) and small intestine (SI) (on the right) of the corresponding boxed regions (on the left) are shown. Scale bar = 250 µm. Arrows show regions of transmural inflammation/crypt distortion, immune infiltrates (in F) correspond also to transmural fibrosis (in H). Segments in between these patches appear normal. **(G)** Bar graphs show the histology index (116) (based on submucosal inflammation, percent area involved, inflammatory infiltrates in LP and crypt hyperplasia and the degree of fibrosis (**I**), as assessed by H&E and trichrome staining on n = 5 WT and 5 GIV-KO mice. *<u>Statistics</u>*: *p* values were determined by unpaired t-test. All results are displayed as mean ± SEM.

Upon *Citrobacter* challenge (**Figure 4C**), GIV-KO mice exhibited an increased acute fecal bacterial load (**Figure 4D**; 1^st^ wk) and an abnormal delay in bacterial clearance, leading to chronic infection (**Figure 4D**; 7^th^ wk). These mice also demonstrated hallmark features of chronic colitis, including colon shortening (**Figure 4E**), patchy transmural inflammation affecting the small intestine, colon, and rectum (**Figure 4, F and G**), as well as focal muscle hypertrophy and collagen deposition (**Figure 4, H and I**). Because the absolute numbers of macrophages and specifically, M2 macrophages—defined by established conventional markers CD68 and CD163, respectively (52–54)—were comparable between *Citrobacter*-infected control and GIV-KO intestinal tissues (**Figure S3, A-D**), we conclude that GIV deficiency impairs the healing functions of ColAMs without affecting macrophage trafficking or polarity-defining M2 markers at the site of infection.

Collectively, these findings highlight a critical role of GIV in bacterial clearance and the resolution of inflammation. Its absence promotes dysbiosis and chronic infectious colitis, underscoring GIV’s essential role in maintaining intestinal immune homeostasis.

### Protective MDP/NOD2 signaling is abolished in myeloid specific GIV-KO mice

Prior studies have shown that pre-treatment with MDP ameliorates infection/bacteremia (55, 56), fatality in sepsis (57) and chemical (e.g., trinitrobenzene sulfonic acid, TNBS and DSS)-induced colitis (15); we asked if these protective actions of MDP require GIV. Compared to WT controls, we found that the GIV-KO mice developed significantly worse DSS-induced acute colitis (**Figure 5A**), as determined by disease activity index (**Figure 5, B and C**) and histological composite scores accounting for deformation of colon crypts and increased immune infiltration in the colon (**Figure 5, D and E**) and; the latter is a composite score of stool consistency, weight loss and the presence of fecal blood (22, 58, 59). Pre-treatment with MDP ameliorated the severity of colitis in WT, but not GIV-KO mice (**Figure 5, B-E**). Because the absolute numbers of CD68+ve M1 and CD163+ve M2 macrophages were comparable between control and GIV-KO DSS-exposed intestinal tissues (**Figure S3E-H**), GIV deficiency appears to impact MDP-induced ColAM properties without affecting macrophage trafficking or polarity-defining M2 markers at the site of inflammation.

**Figure 5.**
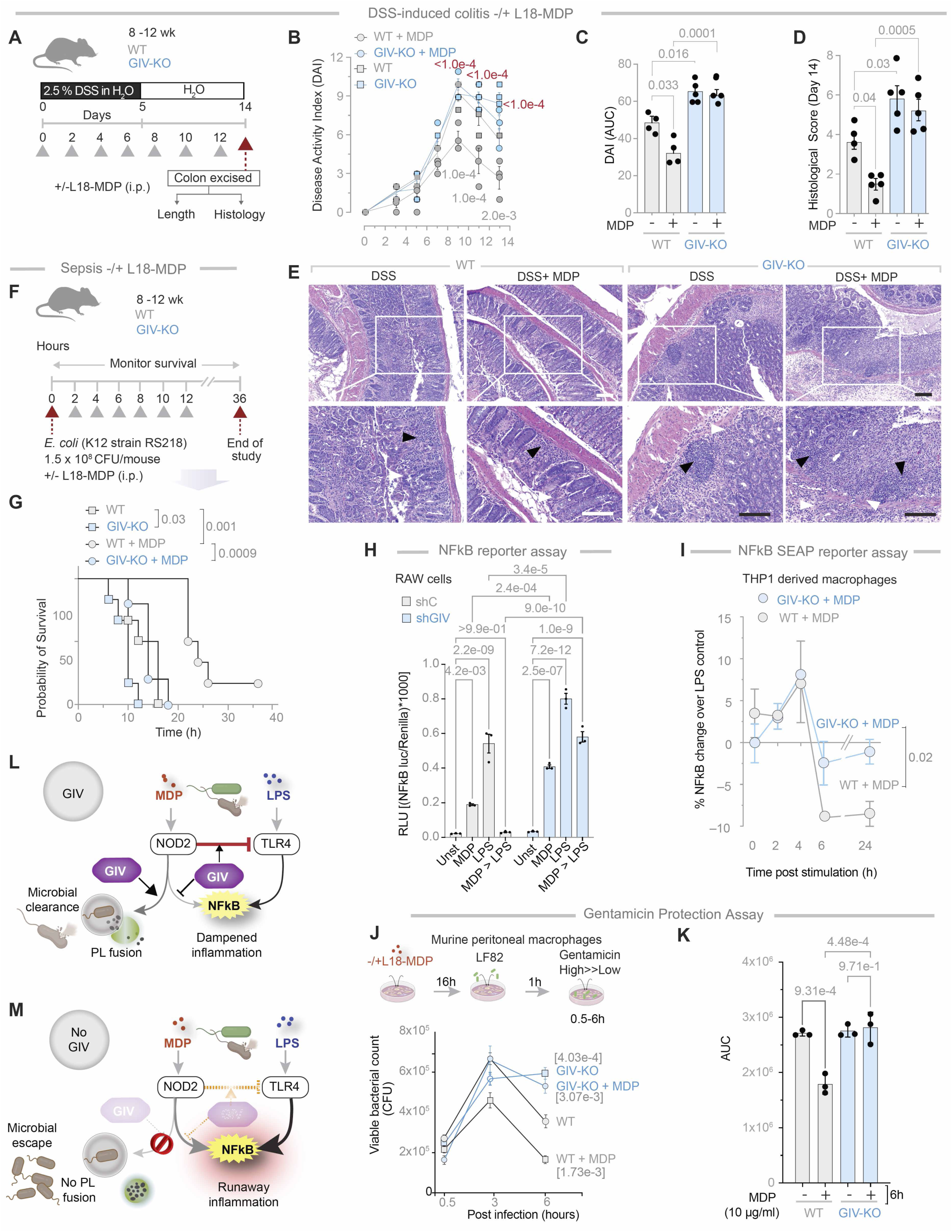
GIV-KO mice and cells are insensitive to the protective actions of MDP/NOD2 signals. **A-E**. Schematic (**A**) displays the study design for DSS-induced colitis. GIV-KO, n=5; WT, n=5. Findings are representative of two independent repeats. Grey arrowheads denote the alternate day administration of muramyl dipeptide (MDP) (100µg/mouse/day). **B-C**. Line graphs (**B**) display disease activity index (DAI), calculated for the days 3, 5, 7, 9, 11 and 13 after DSS administration, which accounts for stool consistency (0-4), rectal bleeding (0-4), and weight loss (0-4). *p* value in gray and red color text represents significance WT vs WT+MDP and WT+MDP vs GIVKO+MDP respectively. Bar graphs (**C**) represent the data in B as area under the curves (AUC) in B. Bar graphs (**D**) display the histological score on day 14, as assessed by a well-accepted methodology (107) of analyzing H&E-stained distal colons from the mice. Representative images are displayed in **E**. Arrowheads point to regions of crypt destruction and/or inflammatory infiltrates. Scale bar = 200 µm. **F-G**. Schematic (A) displays the sepsis study design in which 8 mice in each group were treated with E coli and MDP simultaneously, followed by periodic checks for death (arrowheads). Kaplan-Meier plot (B) displays the % of cohort that survived at those time points. GIV-KO, n=8; WT, n=8. Findings are representative of two independent repeats. *p* values were determined by Mantel-Cox log rank test. *p*-value ≤ 0.05 is considered significant. All results are displayed as mean ± SEM. **H**. Bar graphs display the impact of MDP (10 μg/ml) priming on LPS (100 ng/ml)-induced NFκB activity. **I**. Line graphs display percent change in LPS (100 ng/ml)-induced NFκB activity in WT and GIV-KO cells primed with MDP (10 μg/ml). **J-K**. Schematic (top) displays the experimental setup for bacterial clearance. Line graphs (below) show the viable bacterial counts in the macrophages. Bar graphs (**J**) represent the data in (**I)** as the area under the curve (AUC). **L-M**. Schematic summarizing findings in cells with (**L**) or without (**M**) GIV. *Statistics*: All results are displayed as mean ± SEM (n=3 independent biological replicates). Significance was tested using two-way/one-way ANOVA followed by Tukey’s test for multiple comparisons. *p*-value ≤ 0.05 is considered as significant.

Similar results were observed in the case of *E. coli*-induced sepsis (**Figure 5F**); fatality was higher in GIV-KO mice compared to WT controls (**Figure 5G**). Pre-treatment with MDP reduced fatality in WT, but not GIV-KO mice (**Figure 5G**). These findings demonstrate that GIV is required for the protective MDP/NOD2 signaling in the setting of infection/inflammation.

Prior studies have shown that MDP priming of NOD2 protects cells from excessive inflammation induced by lipopolysaccharide (LPS) (15, 60). To determine if this protective effect requires GIV, we used GIV-depleted (shGIV) RAW 264.7 murine macrophages and WT controls (shC) to assess NFκB activation following LPS stimulation, with or without MDP pretreatment. In WT cells, MDP pretreatment significantly reduced NFκB activation, but this protective effect was markedly compromised in GIV-depleted cells (**Figure 5H**). The findings were also reproduced in CRISPR-depleted human THP-1 cells expressing an NFκB activity-tracking reporter, enabling continuous monitoring of signaling dynamics. The presence of GIV was required for sustained suppression of NFκB activity, evident as early as 6 h and maintained through 24 h (**Figure 5I**). Similar findings were observed in HeLa cells, which express the MD2 co-receptor essential for LPS/TLR4 signaling (48, 61–65) , albeit at low levels (66). In control HeLa cells, MDP pretreatment reduced NFκB activation significantly, both with endogenous NOD2 (**Figure S1F**) and exogenously overexpressed NOD2 (**Figure S1G**). In cells without GIV, this protective effect was either diminished (**Figure S1F**) or virtually abolished (**Figure S1G**). Additionally, when MDP-primed TGPMs were infected with the pathogenic *AIEC*-LF82 strain (49), MDP treatment accelerated bacterial clearance in WT cells but not in KO TGPMs (**Figure 5, J-K**).

These findings demonstrate that GIV is essential for protective MDP/NOD2 signaling, which counteracts LPS/TLR4-driven proinflammatory NFκB signaling (**Figure 5L**). Without GIV, NFκB signaling becomes excessive, and bacterial clearance is delayed and impaired (**Figure 5M**).

### MDP/NOD2 signals induce niColAMs and GIV is required for such induction

To assess the role of MDP/NOD2 signaling in modulating colonic i/niColAM populations, we analyzed publicly available transcriptomic datasets of DSS-induced colitis spanning acute, chronic, and recovery phases (**Figure 6A**), using composite gene signatures of i/niColAM subsets. iColAMs were elevated during the acute phase but declined during the chronic and recovery phases (**Figure 6B**), while niColAMs showed the opposite trend—increased abundance during recovery (**Figure 6B**).

**Figure 6.**
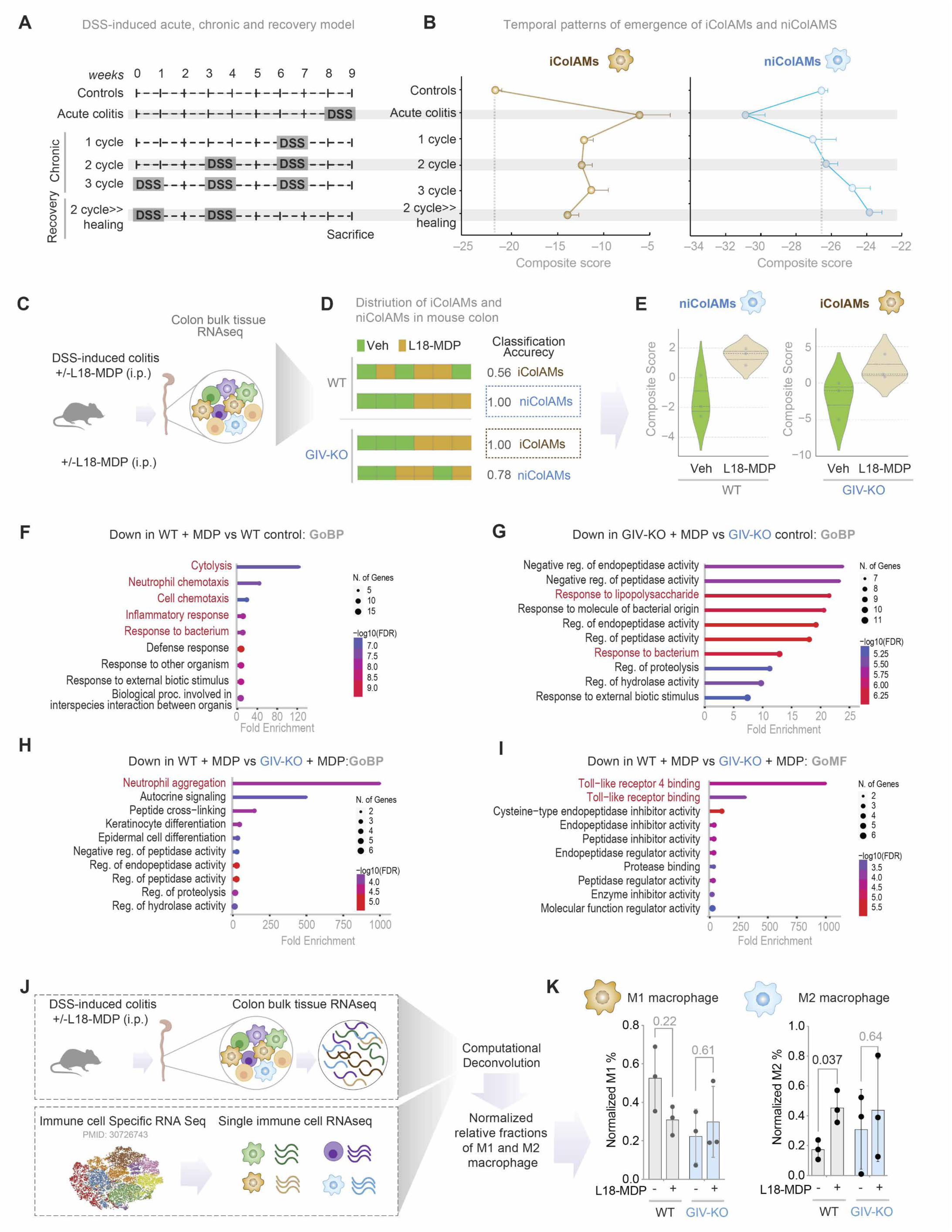
GIV is required for the emergence of healing niColAMs in MDP-treated WT mice. **B.** Study design of DSS-induced acute, chronic and recovery in mouse models of colitis (C57/BL6; all WT). **C.** Line graphs display the temporal patterns of the emergence of iColAMs and niColAMs in the colon samples in A. The grey dotted line indicates the composite scores of iColAM and niColAM genes in the control mice. **C-E.** Study design (**C**) for DSS-induced colitis in WT vs. GIV mice (n=3 each). See also Figure 5A-E for the detailed study design and disease pathology. Bar plots (**D**) show the accuracy of sample classification of individual DSS-challenged mouse samples—treated or not with vehicle (Veh) *vs*. MDP---using iColAM and niColAM gene sets. Classification strength within each cohort is measured using ROC-AUC analyses. Violin plots (**E**) show composite scores for niColAMs (left, blue border) in WT and iColAMs (right, brown border) in GIV-KO mice treated with or without MDP. **F-G**. Gene Ontology Biological Process (Go-BP) pathway enrichment analyses of genes downregulated in WT (F) or GIV-KO (G) mice treated with L18-MDP compared to their respective untreated controls. **H-I**. GoBP (H) and Go Molecular Function (GoMF; I) analyses of genes downregulated in L18-MDP-treated WT vs. GIV-KO samples. **J-K.** Schematic (J) of bulk RNA sequencing *in silico* deconvolution analysis of distal colons from DSS-treated mice in C. Bar plots (K) show normalized percentage abundances of M1 and M2 macrophages in WT and GIV-KO mice, with and without MDP treatment. *Statistics*: *p* values were calculated using one-way ANOVA, followed by Tukey’s test for multiple comparisons and indicated with *p*-values shown above bars. *p*-value ≤ 0.05 is considered significant.

Because GIV is required for the protective effects of MDP/NOD2 signaling in DSS-induced colitis (**Figure 5, A-E**), we examined whether this protection involves modulation of iColAM and niColAM populations in a GIV-dependent manner. We analyzed colon transcriptomes from WT and GIV-KO mice treated with DSS-colitis, with or without MDP treatment (**Figure 6C**). In WT mice, a composite niColAM score robustly distinguished MDP-treated WT colon tissues from untreated WT controls (classification accuracy = 1; **Figure 6D**). This indicates early upregulation of healing niColAMs by MDP—earlier than anticipated from phase-specific trends (**Figure 6B**)—and potentially accelerating recovery. By contrast, in GIV-KO mice, MDP treatment elevated iColAM scores (also with classification accuracy = 1; **Figure 6D**), consistent with the observed exacerbation of inflammatory responses (**Figure 5, A-E**). Gene Ontology (GO) analysis of differentially expressed genes corroborated these findings, revealing activation of protective and reparative programs in MDP-treated WT mice, but not in GIV-KO mice (**Figure 6, F-I**; see **Supplemental information 3** for full genes list).

To determine whether the healing niColAM population aligns with conventional non-inflammatory macrophage (M2) populations, we performed bulk RNA-seq deconvolution. This revealed a close transcriptional resemblance between niColAMs and M2-like (anti-inflammatory) macrophages, which were enriched in MDP-treated WT, but not GIV-KO, mice (**Figure 6, J-K**).

Together, these results underscore the essential role of GIV in enabling MDP/NOD2-mediated protection in vivo—by selectively promoting the emergence of healing niColAMs that counterbalance pro-inflammatory iColAMs and restore tissue homeostasis.

### The GIV●NOD2 interaction is direct and dynamically regulated by MDP

NOD2 typically exists in an inactive, ADP-bound conformation stabilized by intramolecular interactions (67, 68). Upon binding its ligand, MDP, NOD2 undergoes conformational changes that facilitate ADP-to-ATP exchange, self-oligomerization, and downstream signaling (67, 68) (**Figure 7A**).

**Figure 7.**
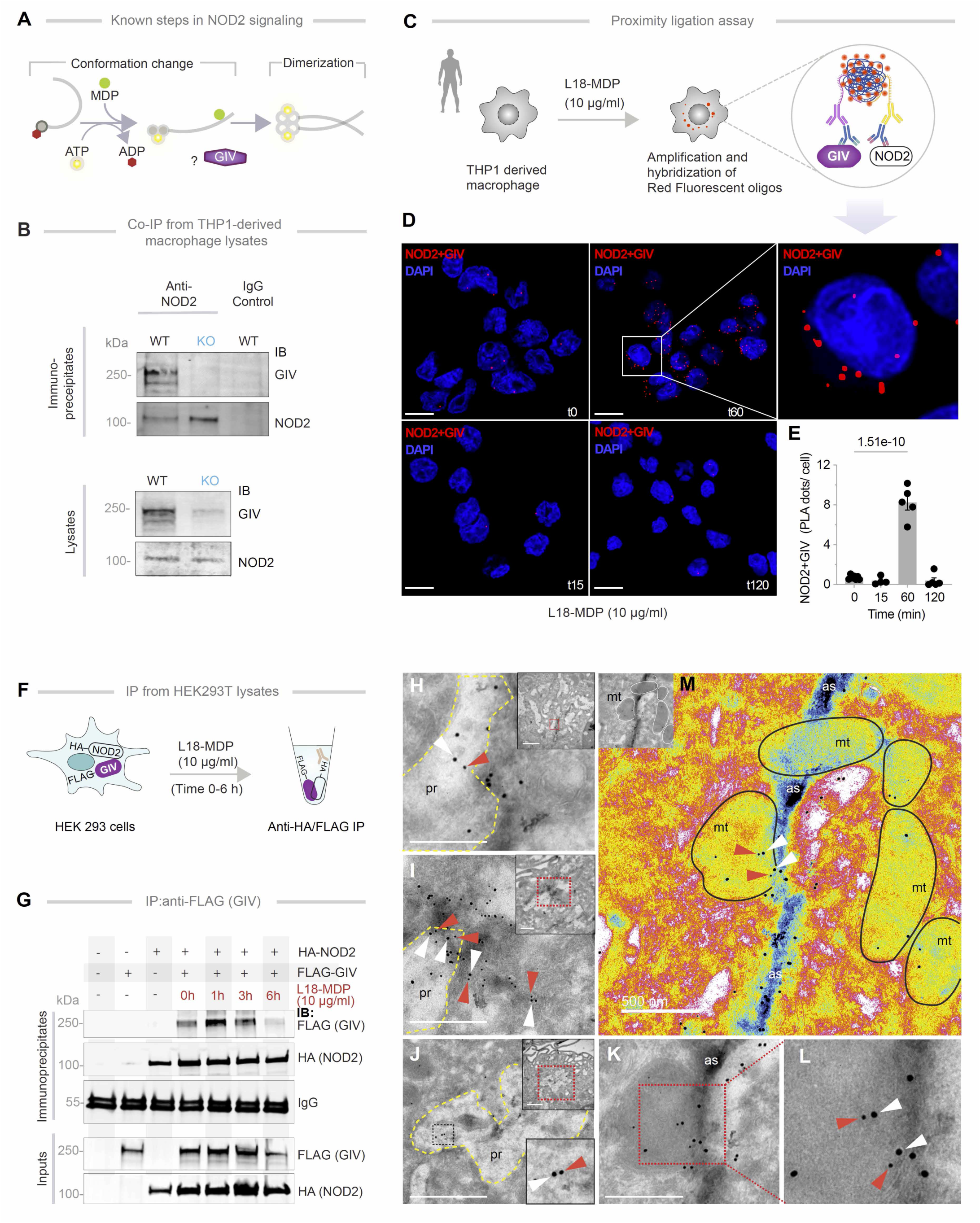
NOD2 and GIV colocalize and interact in cells. **A.** Schematic displays key steps in MDP-induced NOD2 signaling. In resting cells, ADP-bound inactive NOD2 exists in an autoinhibited conformation. Upon ligand (MDP) stimulation, ADP is exchanged for ATP, which further enhances MDP (ligand) binding-associated with conformational change and ‘opening’ of the LRR module. Subsequently, NOD2 dimerizes and forms complexes with other signaling proteins. **B.** Full length endogenous NOD2 was immunoprecipitated from THP1 derived macrophage lysates. Immune complexes were analyzed for bound GIV (top) and equal aliquots of lysates were analyzed for NOD2 and GIV (bottom) by immunoblotting (IB). **C-E**. Study design of PLA (**C**) and representative confocal images (**D**) display colocalization of GIV and NOD2 in THP1 derived macrophages challenged with MDP for 0-120 min. Scale bar = 10 µm. Bar graphs (**E**) display findings based on ∼20-30 randomly imaged fields; n = 4-5 repeats. *p* value was calculated using one-way ANOVA, followed by Tukey’s test for multiple comparisons and indicated with *p*-values shown above bars. *p*-value ≤ 0.05 is considered as significant. **F-G**. Schematic depicts study design of immunoprecipitation from lysates of HEK293T cells (**F**). HA-tagged NOD2 was immunoprecipitated with anti-HA mAb from equal aliquots of lysates of HEK293T cells co-expressing GIV-FLAG and HA-NOD2, stimulated (+) or not (-) with MDP for indicated time points. Immunoprecipitated (IP; top) complexes and input (bottom) lysates were analyzed for NOD2 and GIV by immunoblotting (IB) (**G**). **H-M.** TEM micrographs display representative images of colocalization of GIV (white arrowheads; 18 nm gold particles) and NOD2 (red arrowheads; 12 nm gold particles) on TGPMs challenged with live AIEC*-*LF82 (MOI 1:30) for 1 h. NOD2 colocalization within particle-rich cytoplasmic structures (pr), with membrane-associated GIV on actin strands (as, Pseudo-colored blue) and swollen mitochondria (mt, outlined in black) with degraded cristae in **M** and. Scale bar = 500 nM.

To determine whether GIV physically interacts with NOD2, we performed co-immunoprecipitation experiments and found that full-length, endogenous GIV and NOD2 form complexes in THP1-derived macrophages (**Figure 7B**). *In situ* proximity ligation assay (PLA) using antibodies against the native proteins confirmed this interaction and revealed that the abundance of GIV●NOD2 complexes is enhanced by MDP stimulation, peaking around 1 hour post-treatment (**Figure 7, C**-**E**). Co-immunoprecipitation assays using exogenously expressed, epitope-tagged GIV and NOD2 proteins further validated this interaction and its ligand-dependent dynamic regulation. Whether GIV-FLAG or HA-NOD2 was used as bait, assembled GIV●NOD2 complexes were detected in immune complexes within ∼1-3 h post-MDP stimulation, and declined by ∼6 h (**Figure 7, F-G, Figure S4A**), underscoring the temporally regulated nature of the interaction.

To visualize the ultrastructural context of GIV●NOD2 complex assembly, we performed immunogold electron microscopy on TGPMs 1 h after MDP stimulation. Using 18 nm and 12 nm gold-conjugated antibodies against GIV and NOD2, respectively, we observed NOD2 colocalizing with membrane-associated GIV, predominantly along actin filaments (**Figure 7, H-L**) within particle-rich cytoplasmic structures (69) (PaCS; which contain polyubiquitinated proteins and proteasomes) (**Figure 7, H and J**) and around swollen, morphologically abnormal mitochondria (**Figure 7M**).

Together, these findings demonstrate that the GIV●NOD2 interaction is both direct and dynamically regulated by MDP. The complex associates with the membrane and cytoskeletal elements, supporting its potential role in NOD2-mediated signaling and cellular responses (70).

### The C-terminus of GIV directly binds the LRR domain of NOD2

GIV is a large, multimodular scaffold protein (1871 amino acids) with several defined interaction domains (**Figure 8A**). NOD2, in contrast, contains three major domains: CARD, NBD, and LRR (**Figure 8B**). While NOD1 and NOD2 share structural similarities, co-immunoprecipitation analyses revealed that GIV specifically binds to NOD2 but not NOD1 (**Figure 8C**), indicating that the GIV●NOD2 interaction is selective.

**Figure 8.**
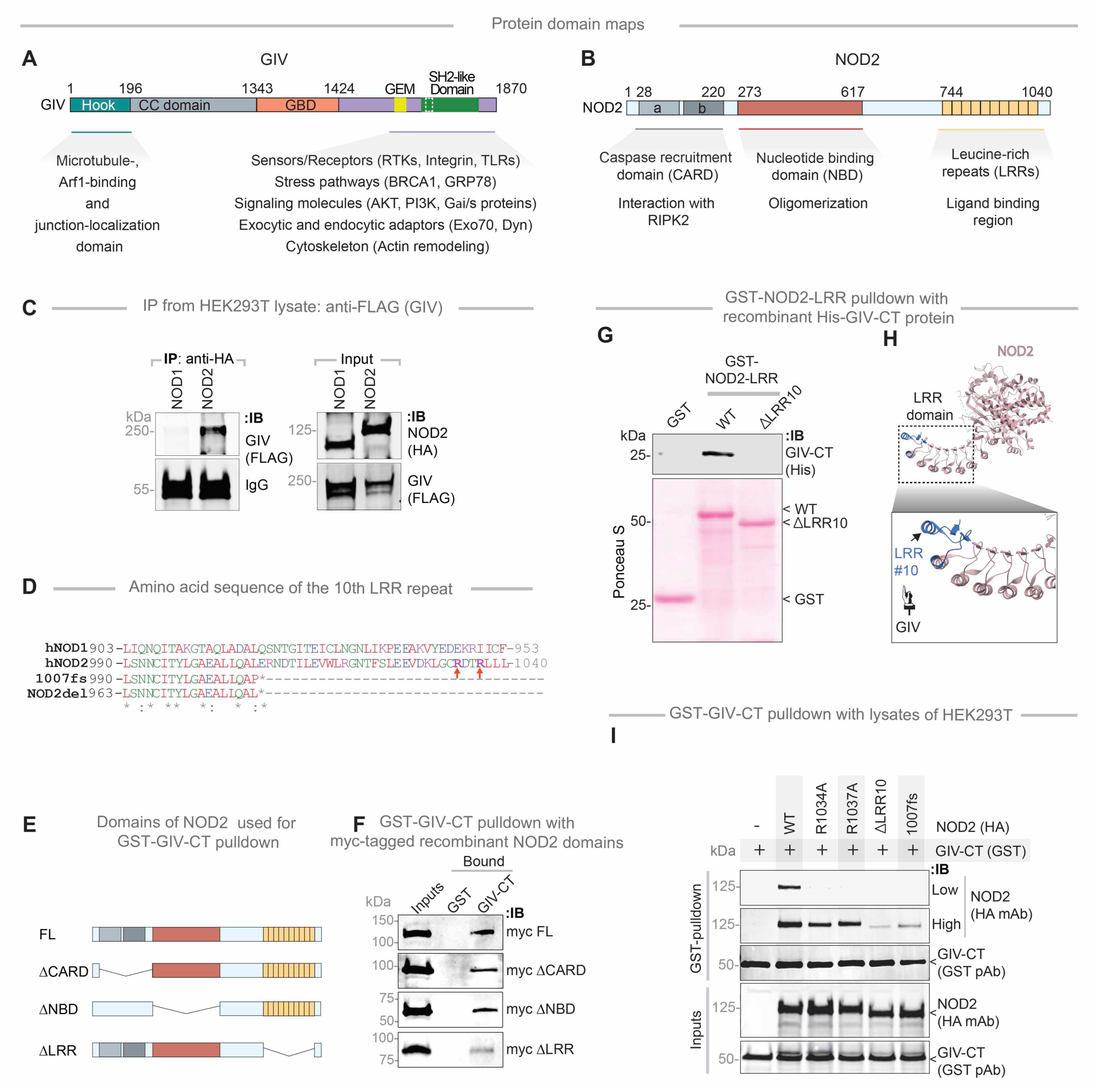
The NOD2(LRR)●GIV(C-term) interaction is direct and ligand-dependent. **A-B.** Schematic depicts the domain maps of (A) GIV and (B) NOD2 and highlights established functions and interactions facilitated by the domains. **C.** HA-tagged NOD1/2 proteins were immunoprecipitated from equal aliquots of lysates of HEK293T cells co-transfected with GIV and either NOD1 or NOD2 using anti-HA mAb. Immunoprecipitated (IP; left) complexes and input (right) lysates were analyzed for NOD1/2 and GIV by immunoblotting (IB). **D.** An alignment of the amino acid sequence of the 10^th^ LRR repeat of human hNOD1, hNOD2, the CD-risk associated NOD2 variant (NOD2-*1007fs*) and the deletion mutant generated in this work (NOD2-del) is shown. Residues mutated in this study to evaluate potential participating residues in the NOD2•GIV interaction are highlighted. **E-F**. Schematics indicate the domains of NOD2 that were used to generate myc-tagged recombinant proteins for use in GST-pulldown assays in **E**. Equal aliquots of recombinant myc-NOD2 domains (∼3 µg; input, **F**) were used in pulldown assays with immobilized GST and GST-GIV in **F**. Myc-tagged NOD2 was visualized by immunoblot (IB) using anti-myc antibody. **G-H**. GST-pulldown assay (G) was carried out using GST NOD2-LRR proteins as indicated and bound His-GIV-CT is assessed. Schematic (H) highlights the terminal LRR repeat (blue) of NOD2 which binds GIV. **I.** GST-GIV-CT was pulled down using Glutathione beads from equal aliquots of lysates of HEK293T lysates co-expressing GST-GIV-CT (aa 1660-1870; mammalian p-CEFL vector) and either WT or HA-NOD2 mutants predicted to disrupt NOD•GIV binding. Immunoprecipitated (IP; top) complexes and input (bottom) lysates were analyzed for NOD2 and GIV-CT by immunoblotting (IB), using anti-HA (NOD2) and anti-GST (GIV-CT) antibodies.

Given that the ∼210 amino acid C-terminal (CT) module of GIV mediates interactions with a variety of receptors and sensors via short linear motifs (SLIMs) (**Figure 8A**), we tested whether this region was sufficient for NOD2 binding. Indeed, GIV-CT bound NOD2 (**Figure S4B**) but did not interact with NOD1 (**Figure S4B**), further confirming specificity.

To identify the domain of NOD2 responsible for GIV binding, we performed co-immunoprecipitation assays using NOD2 deletion mutants lacking the CARD domains (ΔCARD), NBD (ΔNBD), or LRR (ΔLRR) domains (**Figure 8, E and F**). These studies showed that GIV binding was independent of the CARD and NBD domains and instead required the LRR domain (**Figure 8G; Figure S4, C and D**). Notably, deletion of the terminal repeat in the LRR domain (ΔLRR10) virtually abolished GIV binding (**Figure 8, D and G**). These findings were corroborated by co-immunoprecipitation using full-length proteins (**Figure S4E**) and GST-pulldown studies using GIV-CT (**Figure 8I**, **Figure S4F**). Moreover, site-directed mutagenesis of key arginine residues in NOD2 (R1034 and R1037) (**Figure 8D)**, which stabilize its terminal LRR (**Figure 8H)**, significantly reduced GIV binding (**Figure 8I**).

Together, these results demonstrate that the terminal LRR repeat of NOD2 is essential for its interaction with the C-terminal region of GIV, providing insights into the molecular basis of their functional coupling (**Figure 8G-I**).

### GIV fails to bind the CD-associated NOD2 *1007fs* variant, which lacks the terminal LRR

We next examined whether GIV binding is altered by CD-associated NOD2 variants (*R702W*, *G908R* and *1007fs*) which collectively account for ∼80% of mutations associated with CD susceptibility (**Figure 9A**) (32, 71). These mutations affect residues located within or near the LRR domain (**Figure 9B**). Co-immunoprecipitation assays revealed that two variants, R702W and 1007fs (**Figure 9C**), did not bind to the C-terminal region of GIV. Notably, these variants are associated with high disease penetrance (∼100%; **Supplementary Table 1**). By contrast, the G908R variant—which disrupts the MDP-binding interface (72, 73)—retained GIV binding comparable to NOD2-WT (**Figure 9C**). Further studies confirmed that the *1007fs* variant remained incapable of binding GIV even upon MDP stimulation (**Figure 9D**).

**Figure 9:**
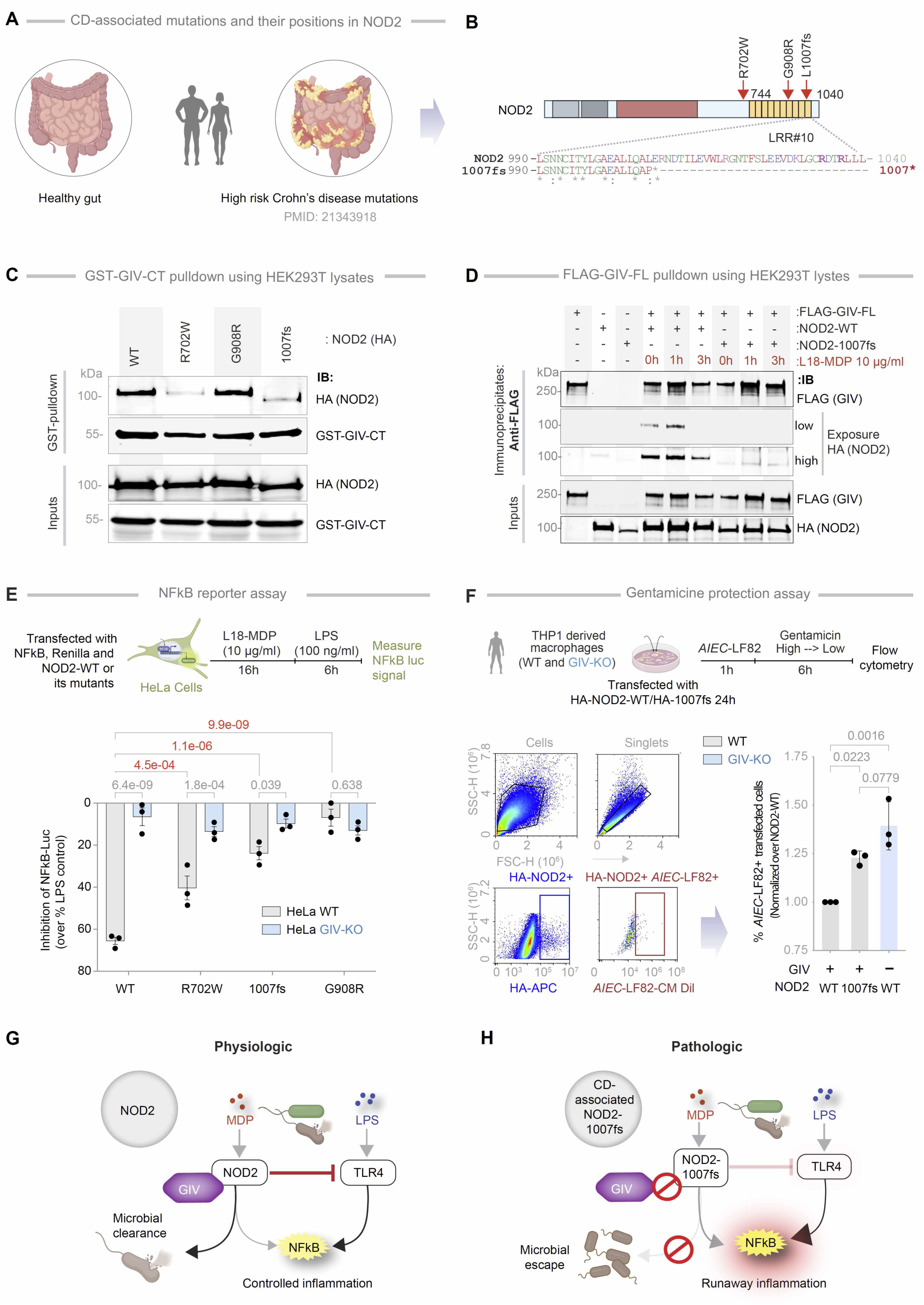
Characterization of the NOD2•GIV interface exploiting CD associated NOD2 mutants. **A-B.** Schematic shows (A) CD-associated mutations and their positions in NOD2, depicts as a domain map (B). *Bottom*: An alignment of the amino acid sequence of the 10^th^ LRR repeat of human NOD2 and the CD-risk associated NOD2 variant (NOD2-*1007fs*) are shown. Residues mutated in this study to evaluate potential participating residues in the NOD2•GIV interaction are highlighted. **D.** GST-GIV-CT was pulled down using Glutathione beads from equal aliquots of lysates of HEK293T co-expressing GST-GIV-CT (aa 1660-1870; mammalian p-CEFL vector) and wild-type (WT) or 3 indicated CD-risk associated variants of HA-NOD2. Bound NOD2 proteins and similar expression of GIV-CT was assessed by immunoblotting (IB) using anti-HA (NOD2) and anti-GST (GIV-CT) antibodies. **E.** FLAG-tagged GIV was immunoprecipitated with anti-FLAG mAb from equal aliquots of lysates of HEK293T cells expressing GIV-FLAG and either wild-type (WT) or *1007fs* variant of HA-NOD2, stimulated (+) or not (-) with MDP for the indicated time points. Immunoprecipitated (IP; top) complexes and input (bottom) lysates were analyzed for NOD2 and GIV by immunoblotting (IB), using anti-HA (NOD2) and anti-FLAG (GIV-CT) antibodies. **F.** Bar graphs display NFκB reporter assay in HeLa cells. Cells were pre-incubated with MDP (10 µg/ml) and then stimulated with LPS (100 ng/ml) and % change of NFκB activity was detected using dual cell reporter assay. **G.** Schematic (top) outlining the workflow for assessing bacterial clearance via flow cytometry. The flow cytometry panel detects CM-Dil labeled *AIEC*-LF82 bacteria (MOI 1:30) in THP1-derived macrophages transfected with HA-NOD2-WT or 1007fs mutant. Bar graphs display *AIEC*-LF82 bacterial load normalized to NOD2-WT. Statistical significance was assessed using one-way ANOVA followed by Tukey’s post hoc test;. *p*-value ≤ 0.05 is considered significant. **G-H**. Schematic summarizing key findings in this work. Magenta colored solid and interrupted lines indicate the GIV-dependent impact on NOD2 that were interrogated in this work. (**F**) *Top*: In physiology, bacterial sensing and signaling by NOD2 requires GIV to limit inflammation. (**G**) In pathology, dysregulated inflammation results when either wild-type NOD2 cannot bind GIV (e.g., GIV is low/absent) or when CD-risk associated *1007fs* variant cannot bind GIV. *Statistics*: *p*-values were calculated using one-way, two-way ANOVA and indicated with *p*-values shown above bars. *p*-value ≤ 0.05 is considered as significant.

To assess the functional consequences of these binding defects, we examined how the CD-risk variants modulate NFκB signaling in response to LPS following MDP priming. In luciferase reporter assays, MDP pretreatment significantly suppressed LPS-induced NFκB activation in the presence of NOD2-WT (**Figure 9E**), conferring 65% protection. However, this protective effect was reduced in cells expressing the NOD2 variants (**Figure 9E**; red *p*-values). For instance, while NOD2-WT conferred ∼65% protection, the NOD2-*1007fs* variant (∼20% protection) or other GIV-binding deficient mutants (**Figure 9E**; red p values). In GIV-KO cells, suppression by NOD2-WT dropped from ∼65% to ∼10%, further confirming the role of GIV in this protective response (**Figure 9E**; gray *p*-values). The residual ∼10–20% suppression observed in conditions lacking GIV●NOD2 coupling (e.g., with NOD2-1007fs or in GIV-KO cells) suggests minor contributions from GIV-independent mechanisms or a consequence of endogenous NOD2.

To investigate the impact of GIV on NOD2-dependent microbial clearance, we performed gentamicin protection assays in THP1-derived macrophages transfected with either NOD2-WT or the 1007fs variant, followed by infection with adherent-invasive E. coli (*AIEC*-LF82) (**Figure 9F**). Macrophages expressing NOD2-WT efficiently cleared bacteria, whereas those with the 1007fs variant showed impaired clearance, reflected by elevated intracellular bacterial burden. Notably, GIV-KO macrophages expressing NOD2-WT displayed a defect similar to GIV proficient control macrophages expressing the NOD2 1007fs variant (**Figure 9F**). These findings indicate that both GIV and the terminal LRR domain of NOD2 (GIV-binding site on NOD2) are critical for effective NOD2-mediated bacterial clearance.

Collectively, these findings highlight the critical role of the GIV●NOD2 interaction in mediating the protective effects of MDP signaling. Disruption of this interaction—such as in patients harboring the 1007fs CD-risk variant—may contribute to exaggerated inflammatory responses (via impaired suppression of LPS-induced NFκB activation) and hinder microbial clearance and restoration of intestinal homeostasis (**Figure 9, G-H)**.

## DISCUSSION

Our study presents three major findings: First, we identified a core gene signature that formally defines colon-associated inflammatory and non-inflammatory macrophage states, in both health and IBD. Within this signature, we established GIV (*CCDC88A* gene) as a critical physical and functional interactor of NOD2, enabling protective and homeostatic NOD2 signaling specifically in non-inflammatory macrophage. This protective mechanism GIV●NOD2 axis operates within the lamina propria across models of acute colitis (DSS-induced), chronic inflammation (IBD), and acute systemic infection (sepsis). Third, we delineated the molecular basis of the GIV–NOD2 interaction, showing that it is direct, dynamic, and essential for dual antimicrobial and anti-inflammatory macrophage responses to bacterial sensing (**Figure 9, G and H**). Specifically, this interaction (i) dampens NFκB-dependent inflammatory signals, and (ii) enhances NFκB-independent pathways that drive phagolysosomal fusion and bacterial clearance. Together, these dual functions prevent excessive inflammation while ensuring effective microbial control.

Importantly, our findings also reveal a molecular mechanism for the pathogenicity of the high-penetrance CD-associated NOD2 variant, 1007fs. In the dysbiotic colitic gut, where NOD2 is essential for regulating inflammation and microbial clearance, the inability of GIV to bind the truncated NOD2-*1007fs* variant provides mechanistic insight into how this risk allele contributes to persistent inflammation, dysbiosis and mucosal pathology. These insights redefine the molecular logic of innate immune sensing and signal integration through NOD2 in intestinal macrophages

### ColAM Signatures Provide a Computational Framework to Map Macrophage States in the Gut

Our machine learning–assisted analyses identified a subset of genes—ColAMs—from a broader macrophage activation signature (SMaRT; 338 genes), which reliably distinguish inflammatory (iColAM) from non-inflammatory (niColAM) macrophages in colon tissue. The ColAM signature, particularly its 53-gene IBD-associated subset, is clinically relevant and reflects dynamic, disease-relevant macrophage states in transcriptomic datasets.

Notably, our findings show that IBD-ColAMs enrich for gut-relevant pathways and successfully resolved macrophage functional states even in bulk RNA-seq datasets, attesting to their robustness and specificity. In fact, we show that i/niColAMs dynamically reflect shifts in macrophage function that track with colitis severity—something conventional markers (like CD163) fail to do.

In healthy tissue, niColAMs predominate, likely reflecting the need for tolerogenic surveillance in a microbe-rich environment protected by a single epithelial layer. These macrophages may act as “brakes”— providing low-grade, tolerogenic surveillance that protects epithelial stem cells and neurons from collateral damage. In contrast, during chronic inflammation and dysbiosis, iColAMs act as “accelerators,” while niColAMs act as “brakes” to restrain runaway inflammation. The niColAMs in the setting of colitis appear to transcriptionally resemble M2 macrophages, which have been implicated in mounting an adequate healing response. The niColAMs are induced by NOD2 activation and GIV appears to be essential for such induction. Disruption of this balance—whether through hyperactive iColAMs or impaired niColAMs (as seen in GIV KO mice)—may perpetuate inflammation and disease. This “brake and accelerator” framework offers a new conceptual framework for understanding macrophage regulation at mucosal barriers and presents a foundation for therapeutic targeting of macrophage states in IBD.

### GIV Enables NOD2 to Restrain NFκB-Driven Inflammation in Non-Inflammatory Macrophages

Among the 53 IBD-ColAM genes, *CCDC88A* (GIV) emerges as the sole candidate that both physically and functionally interacts with NOD2. While NOD2’s suppression of NFκB-driven inflammation is well recognized but poorly understood (13), we now show that GIV is essential for this protective function. GIV physically interacts with NOD2, and such binding enables NOD2 to (i) suppress excessive NFκB activity and (ii) drive bacterial clearance via cytoskeletal and phagolysosomal pathways—both essential for mucosal immunity and homeostasis. In the absence of GIV, NOD2’s antimicrobial and anti-inflammatory functions are impaired, macrophages adopt a reactive phenotype, and host defenses falter. Consequently, macrophages adopt reactive phenotypes, display impaired microbial control, and the host shows heightened susceptibility to colitis and sepsis. Notably, GIV’s inability to bind the NOD2-1007fs variant supports a molecular mechanism linking GIV to chronic intestinal inflammation. These data position GIV as a central integrator of gut immune regulation and tissue repair.

These findings build on prior work showing GIV acts as a “brake” within the LPS/TLR4 signaling cascade (22). GIV’s conserved C-terminal motif binds and dampens inflammatory signaling by TLR1/2, TLR2/6, and TLR3, inducing tolerogenic programs aimed at homeostasis and immunity (22). Thus, GIV emerges as a point of convergence for major pattern recognition receptors (PRRs), coordinating tolerogenic responses during microbial sensing (22, 37, 74).

### GIV Couples NOD2 to Other NFkB-independent Signaling Domains and Organelle Functions

Our mechanistic analyses show that GIV binds NOD2 via its C-terminal 210 amino acids, interacting specifically with the terminal LRR repeat of NOD2. This identifies GIV as only the third known protein to directly engage the NOD2-LRR region (75), and one of very few to do so in a way that enhances NOD2’s protective, NFκB-suppressive signaling. While our study defines how GIV shapes NOD2 function, the possibility of reciprocal regulation remains unexplored. It is possible that the GIV●NOD2 interaction may collaborate or compete with GIV-dependent cAMP inhibition (via Gi activation and Gs inhibition) (76), or temporally-spatially cross-regulate each other, impacting myriad of inflammatory signals that are shaped by cAMP flux (77–79), including phagolysosomal fusion events that are critical for microbial clearance (80). This would position NOD2 as another receptor modulating trimeric G-proteins and cAMP through GIV’s C-terminal SLIM motifs, joining a lengthy list of priors (81). Future studies will identify the specific SLIM mediating this interaction and investigate overlap with motifs for TLR4 or Gαi/s binding.

Taken together with its impact on NFkB-driven inflammation, the NOD2-GIV module likely evolved to balance pathogen elimination with inflammatory restraint. This dual functionality—dampening NF-κB while ensuring efficient phagolysosomal fusion—may be key to preventing collateral tissue damage during infection and preserving mucosal homeostasis.

### The Loss of GIV●NOD2 Interaction Defines the Functional Defect in the 1007fs CD-Associated Variant

Among the three main CD-associated variants (*R702W*, *G908R*, and *1007fs*) that interfere with bacterial recognition (82), G908R’s defect lies in impaired MDP contact (72, 73) whereas R702W and 1007fs show defects in palmitoylation and PM localization (37). Only the *1007fs* variant, which lacks the terminal LRR repeat, fails to regain functionality upon restoring PM localization (83), indicating that *1007fs* variant lacks key functions, perhaps because of the truncated terminal LRR repeat. We show that the *1007fs* variant does not bind GIV and that it lacks the same terminal LRR repeat that is essential for the NOD2●GIV interaction. Because our conclusions are supported by both co-immunoprecipitation and *in vitro* pulldown assays using recombinant NOD2-LRR proteins, it unlikely that mislocalization artifacts explain the binding loss [as proposed for other NOD2 interactors, e.g., Erbin (84)]. Thus, GIV emerges as a first-in-class NOD2-interactor that specifically requires the terminal (10^th^) LRR repeat —precisely the region lost in the 1007fs variant. The observation that NOD2-1007fs–expressing cells phenocopy GIV-deficient cells, exhibiting heightened inflammation and impaired microbial clearance, further underscores the critical role of this terminal repeat as the GIV-binding site, whose loss disrupts NOD2’s protective signaling.

Because this variant (also termed *3020insC*) is most consistently associated with CD across multiple studies and population groups, and displays 100% disease penetrance, it is not surprising that our GIV-KO animals challenged with *Citrobacter* develop key features of CD : patchy ileocolitis, transmural inflammation, focal muscle hypertrophy, fibrosis, and dysbiosis. Notably, these features arise within just 7 wks — substantially earlier than the only other known spontaneous murine model of CD SAMP1/YitFcs, which takes ∼30 wks (85). Future studies will explore whether GIV-KO mice recapitulate the full molecular and phenotypic spectrum of Crohn’s disease, including defective innate/adaptive immunity and fistula formation.

## LIMITATIONS OF STUDY

Although our conclusions are grounded in NOD2-specific phenotypes elicited by MDP stimulation, we lacked tools to directly interrogate the GIV●NOD2 interaction in vivo. Future studies will require engineered mutants of GIV and NOD2 that selectively disrupt binding, enabling direct assessment of interaction-dependent functions. Additionally, the observation that MDP enhances GIV●NOD2 binding raises the possibility that other variables—such as pH, ATP levels, or subcellular localization—may modulate this interaction. These contextual factors, known to influence the NOD2 interactome, were not investigated in this study but remain important avenues for future exploration. Finally, we know that GIV can modulate signaling downstream of multiple TLRs, and NOD2 can suppress a subset of those TLRs (15, 17, 22, 86). While this study establishes the role of a functional coupling between GIV and NOD2 in dampening TLR4-driven inflammation, further studies are needed to determine whether GIV-dependent NOD2 signaling broadly suppresses TLR-mediated responses beyond TLR4.

## METHODS

### Computational

#### StepMiner analysis

StepMiner is an algorithm that identifies step-wise transitions using step function in time-series data (87). StepMiner undergoes an adaptive regression scheme to verify the best possible up and down steps based on sum-of-square errors. The steps are placed between time points at the sharpest change between expression levels, which gives us the information about timing of the gene expression-switching event. To fit a step function, the algorithm evaluates all possible steps for each position and computes the average of the values on both sides of a step for the constant segments. An adaptive regression scheme is used that chooses the step positions that minimize the square error with the fitted data. Finally, a regression test statistic is computed as follows:

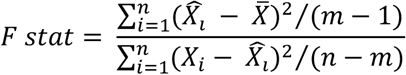

Where *𝑋_i_* for 𝑖 = 1 to 𝑛 are the values, 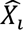 for 𝑖 = 1 to 𝑛 are fitted values. m is the degrees of freedom used for the adaptive regression analysis. *X̄*, is the average of all the values: 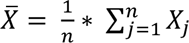. For a step position at k, the fitted values 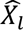 are computed by using 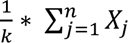 for 𝑖 = 1 to 𝑘 and 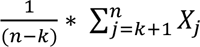 for 𝑖 = 𝑘 + 1 to 𝑛.

#### Measurement of classification strength or prediction accuracy

To measure the strength of classification or prediction accuracy, Receiver Operating Characteristic (ROC) curves were generated for each gene. These curves illustrate the diagnostic ability of a binary classifier system (e.g., high vs. low StepMiner normalized gene expression levels) as its discrimination threshold is adjusted along with the sample order. ROC curves plot the True Positive Rate (TPR) against the False Positive Rate (FPR) at various threshold settings. The Area Under the Curve (AUC) quantifies the probability that a classifier will correctly rank randomly chosen samples into two groups of healthy and IBD patients. Alongside ROC AUC, other classification metrics such as accuracy ((TP + TN)/N; TP: True Positive; TN: True Negative; N: Total Number), precision (TP/(TP+FP); FP: False Positive), recall (TP/(TP+FN); FN: False Negative), and f1 score (2 * (precision * recall)/(precision + recall)) were computed. The Python Scikit-learn package was used to calculate the ROC-AUC values.

#### Composite gene signature analysis using Boolean Network Explorer (BoNE)

Boolean network explorer (BoNE) (7) provides an integrated platform for the construction, visualization and querying of a gene expression signature underlying a disease or a biological process in three steps: First, the expression levels of all genes in these datasets were converted to binary values (high or low) using the StepMiner algorithm. Second, Gene expression values were normalized according to a modified Z-score approach centered around *StepMiner* threshold (formula = (expr - SThr)/3*stddev). Third, the normalized expression values for every gene were added together to create the final composite score for the gene signature. As a modified Z-score, the composite score of a gene signature ranges from negative to positive values, reflecting the dynamic range of each gene in the signature. These composite scores represent the overall activity or state of biological pathways associated with the genes and, hence, can identify differences between control and query groups *within* any given dataset. However, composite scores cannot be directly compared between different gene signatures within a dataset (as they are not normalized according to the number of genes in each signature), nor can the same signature be compared across datasets (which are individually normalized based on intra-dataset sample distribution or have other inherent differences). The samples were ordered based on the final signature score. Differentially expressed genes were identified using DESeq2 R package. Welch’s Two Sample t-test (unpaired, unequal variance (equal_var=False), and unequal sample size) parameters were used to compare the differential signature score in different sample categories. Violin, swarm and bubble plots are created using python seaborn package version 0.10.1. Pathway enrichment analyses for genes were carried out using the KEGG database (88). Violin plots are created using python seaborn package version 0.10.1.

#### Correlation Heatmap

The correlation coefficient between two gene expression values in a transcriptomics dataset was calculated by plotting them in a scatter plot and measuring the correlation coefficient using Python’s SciPy library.

#### Bulk RNAseq Deconvolution

The *in-silico* deconvolution of bulk RNA sequencing data to estimate immune cell-type abundance in murine colon samples was performed using the Granulator R package (89). To normalize cell-type abundances, we utilized the immune cell signature matrix developed by Monaco et al., (90).

### Experimental

#### Cell culture

Thioglycolate-elicited murine peritoneal macrophages (TGPMs) were collected from peritoneal lavage of 8- to 12-wk-old C57BL/6 mice with ice cold RPMI (10 ml per mouse) 4 days after intraperitoneal injection of 3 ml of aged, sterile 3% thioglycolate broth (BD Difco, USA) and cultured as described previously (91). Cells were passed through 70 µm filter to remove possible tissue debris contamination during harvesting. Cells were counted, centrifuged, and resuspended in RPMI-1640 containing 10 % FBS and 1% penicillin/streptomycin. Cells were plated with required cell density and the media was changed after 4 h to remove non adherent cells. Cells were allowed to adjust to overnight culture before the addition of stimuli as indicated in the Figure legends. RAW264.7, HEK293T and HeLa cells (from ATCC) were maintained in the DMEM media containing 10% FBS and 1% penicillin/streptomycin. THP1 reporter cells (THP1-Dual™ Cells, InVivoGen, USA) derived from THP-1, a human monocytic cell line and parental THP1 WT cells were maintained RPMI media containing 10% FBS and 1% penicillin/streptomycin.

#### Bacteria and bacterial culture

*Citrobacter rodentium* (strain DBS100) and Adherent Invasive *Escherichia coli* strain LF82 (AIEC-LF82) were cultured from a single colony inoculation into LB broth for 6-8 h on shaking incubator, followed by overnight culture under oxygen-limiting conditions, without shaking, to maintain their pathogenicity as done previously (92–94). Bacterial cells were counted by measuring an absorbance at 600 nm (OD600), washed with PBS, and infected with indicated MOI in Figure legends.

#### Mice

*Ccdc88a^fl/fl^* mice were a gift from Dr. Masahide Takahashi (Nagoya University, Japan) and was developed as described (95). LysM*^Cre/Cre^* mice (B6.129P2-Lyz2t (90)(cre)lfo/j) were purchased from the Jackson Laboratory. *Ccdc88a^fl/fl^* x LysM*^Cre/-^* mice were generated previously by us as described (22) and were maintained as homozygous floxed and heterozygous LysMcre. Primers required for genotyping are mentioned in Table 1. Both male and female mice (8-12 weeks) were used and maintained in an institutional animal care at the University of California San Diego animal facility on a 12-hour/12-hour light/dark cycle (humidity 30–70% and room temperature controlled between 68–75 °F) with free access to normal chow food and water. All mice studies were approved by the University of California, San Diego Institutional Animal Care and Use Committee (IACUC).

**Table 1:**
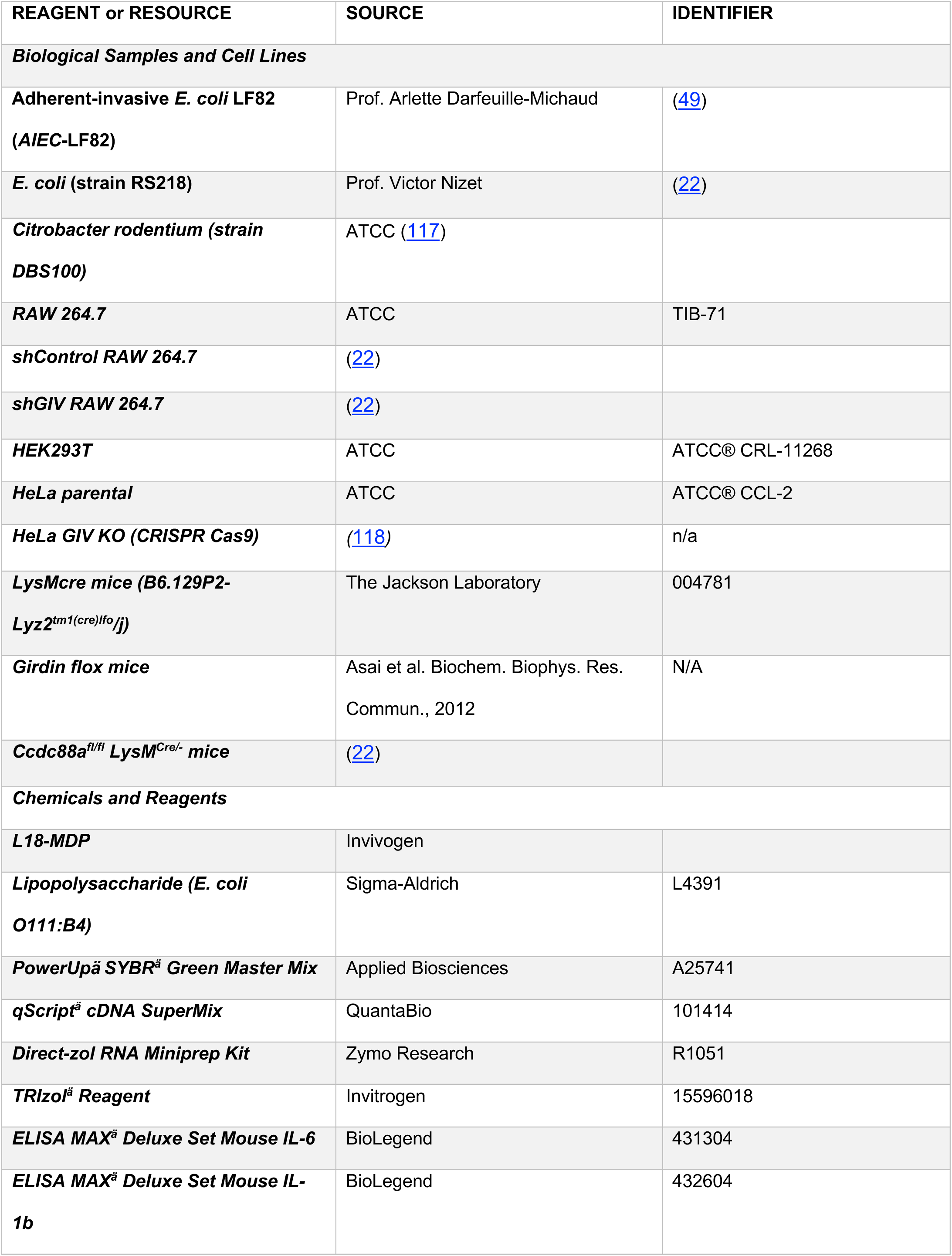
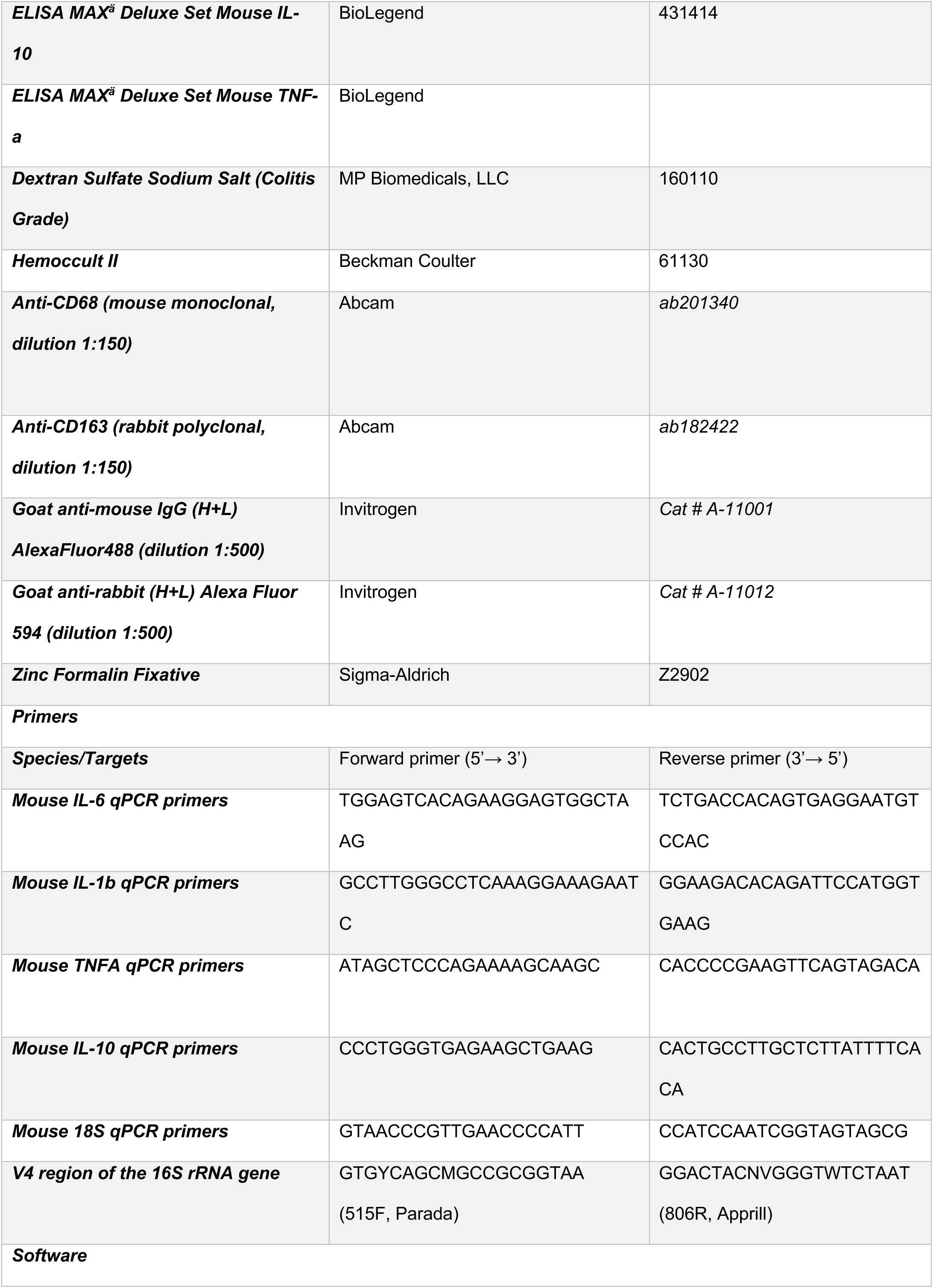
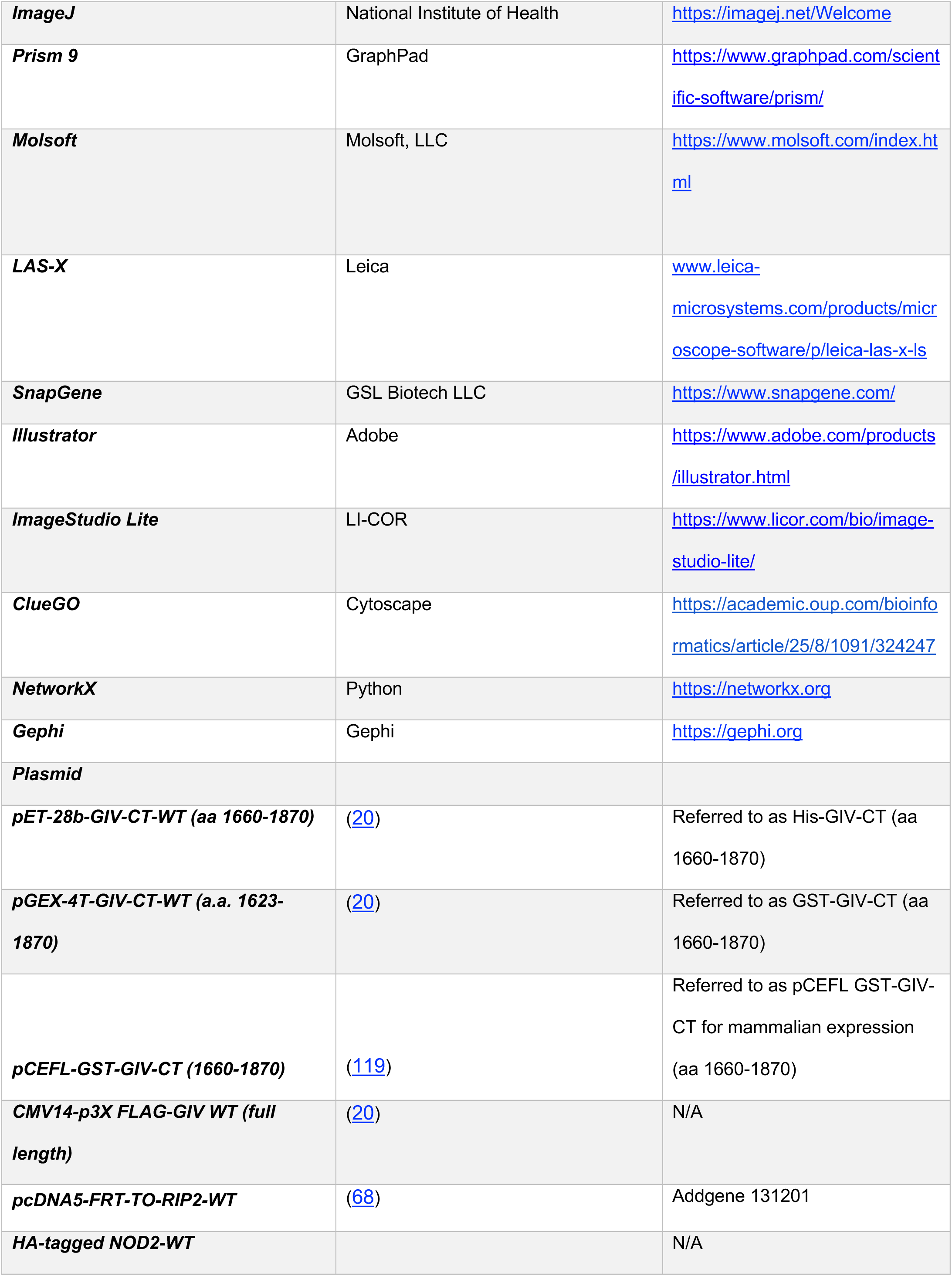
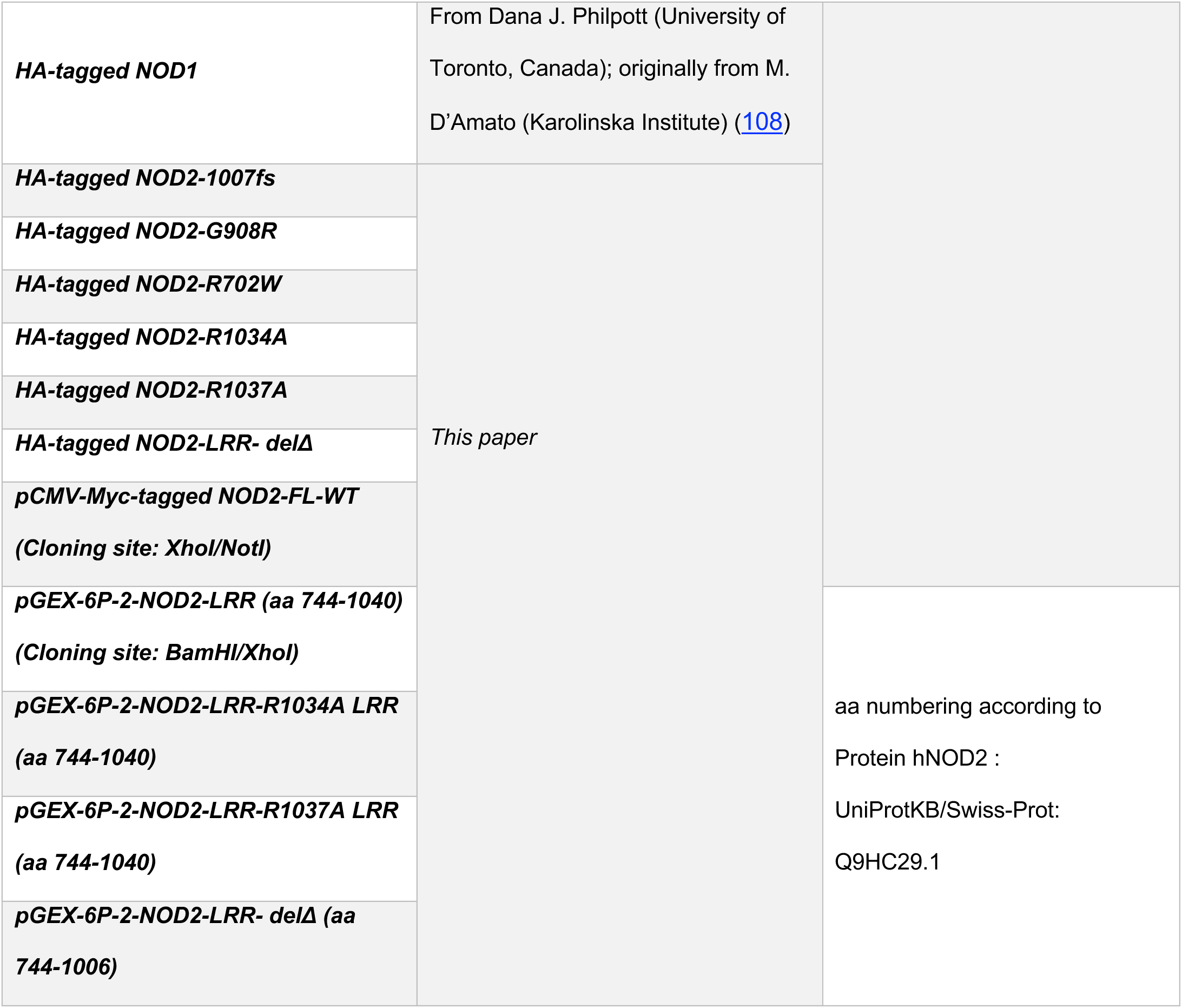
List of reagents and resources.

#### Fecal pellet collection, DNA extraction and 16S rRNA sequencing

Individual mice fecal pellets from co-housed GIV-KO mice and their littermate WT controls (8-12 weeks) were collected in clean containers and frozen at -80°C untill use. Samples were transported to microbiome core facility, University of California, San Diego for 16S rRNA processing. Total DNA was extracted from the individual mice fecal pellets using MagMAX Microbiome Ultra Nucleic Acid Isolation kit, (Thermo Fisher Scientific, USA) and automated on KingFisher Flex robots (Thermo Fisher Scientific, USA). 16S rRNA gene amplification was performed according to the Earth Microbiome Project protocol (96). Briefly, Illumina primers with unique forward primer barcodes (97) were used to amplify the V4 region of the 16S rRNA gene (515F-806R (98)) with single reactions per sample (99). Equal volumes of each amplicon were pooled, and the library was sequenced on the Illumina MiSeq sequencing platform with paired-end 150 bp cycles.

#### 16S rRNA gene data analysis

QIIME2 (100) was used to process the demultiplexed files. Sequences were filtered, denoised and trimmed to 150 bp using DADA2 (101), which is used to correct Illumina-sequenced amplicon errors. The sequences were classified using the q2-feature-classifier plugin from QIIME2 that was trained on the Green genes 13_5 99% OTUs trimmed to 150 bp. Alpha and beta diversity plots were generated using the R microbiome package (102). Alpha diversity was estimated using Faith’s phylogenetic diversity and Shannon diversity. Beta diversity was estimated using Non-Metric Multidimensional Scaling (NMDS) and Bray-Curtis dissimilarity (102).

#### *C. rodentium* induced infectious colitis

*C. rodentium* (strain DBS100) induced infectious colitis studies were performed on 8-week*-old* GIV-KO and their littermate WT control mice. *C. rodentium* were grown overnight in LB broth with shaking at 37 °C. Mice were gavaged orally with 5 x 10^8^ CFU in 0.1 ml of PBS (103, 104). To determine viable bacterial numbers in faeces, fecal pellets were collected from individual mice, homogenized in ice-cold PBS, serially diluted, and plated on MacConkey agar plates. Number of CFU was determined after overnight incubation at 37 °C. Colon samples were collected to assess histology in the 7^th^ week.

#### DSS-induced colitis

To induce colitis, mice were fed with drinking water containing 2.5% dextran sulfate sodium (DSS, w/v) (MP Biomedicals, MW 36–50 kDa) for five days and then replaced normal drinking water as described (105, 106). For treatment study, MDP (100 mg/mouse/day) was administered via intraperitoneal route in 100 ml total volume sterile saline every alternate day starting from day 0 of experiment. Mice were sacrificed on the 14^th^ day, and colon length was assessed. Colon samples were collected to assess the levels of mRNA (by qPCR). Drinking water levels were monitored to determine the volume of water consumption. Weight loss, stool consistency, and fecal blood were recorded fro individual animals, and these parameters were used to calculate an average Disease Activity Index (DAI) as described previously (107). Colon histology was assessed in hematoxylin and eosin-stained tissue sections using standard protocols.

#### *E. coli*-induced sepsis

*E. coli* (strain RS218) was grown overnight in Lysogeny broth (LB) media with shaking at 37°C. Next morning, fresh LB media was used to dilute cultures to 1:50 and grow up to mid-log phase, washed twice with PBS, and reconstituted in PBS. GIV-KO mice (8–12-week-old) and their control WT littermates were injected with 1.5 x 10^8^ CFU of *E. coli* in 200 ul and mice survival was recorded for 24h post-infection. Mice were pre-treated with MDP (100 μg i.p.) or PBS 18 h before the infection.

#### Transmission electron microscopy (TEM) and immunogold EM staining

Cells were fixed with Glutaraldehyde in 0.1M Sodium Cacodylate Buffer, (pH 7.4) and post-fixed with 1% OsO4 in 0.1 M cacodylate buffer for 1 hr on ice. The cells were stained with 2% uranyl acetate for 1 hr on ice and dehydrated in graded series of ethanol (50-100%) while remaining on ice. The cells were washed once with 100% ethanol and twice with acetone (10 min each) and embedded with Durcupan. Sections were cut at 60 nm on a Leica UCT ultramicrotome and picked up on 300 mesh copper grids. All sections were post-stained sequentially 5 min with 2% uranyl acetate and 1 min with Sato’s lead stain. Samples were visualised using JEOL 1400 plus equipped with a bottom-mount Gatan OneView (4k x 4k) camera. For immunogold staining to determine co-localization of NOD2 and GIV. The sections were incubated with mouse anti-NOD2 antibody (Santa Cruz, sc-56168, 1:50 dilution) and rabbit anti-GIV antibody (Millipore sigma, ABT80; 1:50 dilution) followed by secondary antibodies 18 nm colloidal gold of donkey anti-rabbit IgG and 12 nm gold of donkey anti-mouse IgG (Jackson ImmunoResearch Laboratories, Inc.). Samples were visualised using JEOL 1400 plus equipped with a bottom-mount Gatan OneView (4k x 4k) camera.

#### Plasmid constructs

The plasmids used in this study were HA-Nod1 and HA-Nod2 (from Dana Philpott (108) and M. D’Amato (109)); and His-Myc tagged NOD2 constructs (His-myc-Nod2 FL, ΔCARD-His-myc-Nod2, ΔNBD-His-myc-Nod2, ΔLRR-His-myc-Nod2) were generous gift from Santanu Bose (110), . All other Nod2 mutants were generated by site directed mutagenesis. GST-Nod2-LRR and Myc-Nod2 were cloned into pGEX-6P and pCMV-myc vectors, respectively, using cloning sites indicated in Table 1.

#### Protein expression and purification

GST and His-tagged proteins were expressed in *E. coli* strain BL21 (DE3) and proteins were purified as described (22, 111, 112). Briefly, to stimulate protein expression, bacteria cultures were activated with 1 mM IPTG overnight at 25°C. Bacteria were then pelleted and resuspended in either GST lysis buffer (25 mM Tris-HCL (pH 7.4), 20 mM NaCl, 1 mM EDTA, 20% (vol/vol) glycerol, 1% (vol/vol) Triton X-100, protease inhibitor cocktail) or His lysis buffer (50 mM NaH2PO4 (pH7.4), 300 mM NaCl, 10 mM imidazole, 1% (vol/vol) Triton-X-100, protease inhibitor cocktail), sonicated and lysates were cleared by centrifugation at 12,000 x g at 4°C for 30 mins. Supernatant was then affinity purified using glutathione-Sepharose 4B beads or HisPur Cobalt Resin, followed by elution, overnight dialysis in PBS, and then stored at -80°C until use.

#### Transfection, lysis, and quantitative immunoblotting

HEK293T and HeLa cells were cultured in DMEM media containing 10% FBS and antibiotics according to the ATCC guidelines. Cells were transfected using polyethylenimine for DNA plasmids following the manufacturers’ protocols. Lysates for immunoprecipitation assays were prepared by resuspending cells in lysis buffer (20 mM HEPES, pH 7.2, 5 mM Mg-acetate, 125 mM K-acetate, 0.4% Triton X-100, 1 mM DTT) supplemented with 500 µM sodium orthovanadate, phosphatase inhibitor cocktails (Sigma) and protease inhibitor cocktails (Roche), and cleared (10,000 x *g* for 10 min) before use. For immunoblotting, proteins were fractionated by SDS-PAGE and transferred to PVDF membranes (Millipore). Membranes were blocked with 5% nonfat milk dissolved in PBS before incubation with primary antibodies followed by detection with secondary antibodies using infrared imaging with two-color detection and quantification were performed using a Li-Cor Odyssey imaging system. All Odyssey images were processed using Image J software (NIH) and assembled for presentation using Photoshop and Illustrator software (Adobe).

#### Immunoprecipitation, *in vitro* GST-pulldown with recombinant purified protein or cell lysates

For *in vitro* pulldown assays, purified GST-tagged proteins from *E. coli* were immobilized onto glutathione Sepharose beads by incubating with binding buffer (50 mM Tris-HCl (pH 7.4), 100 mM NaCl, 0.4% (vol/vol) Nonidet P-40, 10 mM MgCl2, 5 mM EDTA, 2 mM DTT) overnight at 4ᴼC with continuous rotation. GST-protein bound beads were washed and incubated with purified His-tagged proteins resuspended in binding buffer or with pre-cleared (by centrifugation at 10,000xg for 10 min) cell lysates prepared using lysis buffer [20 mM HEPES, pH 7.2, 5 mM Mg-acetate, 125 mM K-acetate, 0.4% Triton X-100, 1 mM DTT, 500 μM sodium orthovanadate supplemented with phosphatase inhibitor cocktail (Sigma Aldrich) and protease inhibitor cocktail (Roche)] for 4 hrs at 4°C. After binding, bound complexes were washed four times with 1 ml phosphate wash buffer (4.3 mM Na2 HPO4, 1.4 mM KH2PO4 (pH 7.4), 137 mM NaCl, 2.7 mM KCl, 0.1% (vol/vol) Tween-20, 10 mM MgCl2, 5 mM EDTA, 2 mM DTT, 0.5 mM sodium orthovanadate) and eluted by boiling in Laemmli buffer (5% SDS, 156 mM Tris-Base, 25% glycerol, 0.025% bromophenol blue, 25% β-mercaptoethanol).

#### Co-immunoprecipitation assays

N-terminal HA- or Myc-tagged NOD2 and C-terminal FLAG-tagged GIV full length proteins were co-expressed in HEK293T cells and 48 h after transfection, cells were stimulated with 10 µg/ml MDP (L18-MDP, Invivogen) for 1, 3 and 6 h followed by cell lysis in lysis buffer (20 mM HEPES, pH 7.2, 5 mM Mg-acetate, 125 mM K-acetate, 0.4% Triton X-100, 1 mM DTT, 0.5 mM sodium orthovanadate, Tyr phosphatase inhibitor cocktail, Ser/Thr phosphatase inhibitor cocktail, and protease inhibitor cocktail). For immunoprecipitation, equal aliquots of clarified cell lysates were incubated for 3h at 4°C with 2 μg of appropriate antibody [either anti-HA mAb or anti-FLAG M2Ab (Sigma Monoclonal ANTI-FLAG® M2, Clone M2)]. Subsequently, protein G Sepharose beads (GE Healthcare; 40 µl 50% v:v slurry) were added and incubated at 4°C for an additional 60 min. Beads were washed 4 times (1 ml volume each wash) in PBS-T buffer [4.3 mM Na2HPO4, 1.4 mM KH2PO4, pH 7.4, 137 mM NaCl, 2.7 mM KCl, 0.1% (v:v) Tween 20, 10 mM MgCl2, 5 mM EDTA, 2 mM DTT, 0.5 mM sodium orthovanadate] and immune complexes were eluted by boiling in Laemmli’s sample buffer. Bound immune complexes were separated on SDS PAGE and analyzed by immunoblotting with anti-HA, anti-Myc and anti-FLAG antibodies. In assays where the impact of nucleotides was studied on protein-protein interactions, cells were were first permeabilized with 4 μg/ml of streptolysin-O (Sigma-Aldrich, Budapest, Hungary) for 30 min at 37°C followed by incubation with nucleotides (ADP, ATP, or ATPgS).

#### Proximity ligation assay (PLA)

THP1 cells grown on glass coverslips nearly 70% confluency, washed two times with PBS, and fixed 20 min at room temperature with 4% (wt/vol) PFA. After washing with PBS, cells were processed for PLA using the Duolink *In Situ* PLA kit according to the manufacturer’s instructions. Briefly, coverslips were blocked for 1 hour at room temperature, followed by a 1-hour incubation with primary antibodies targeting GIV and NOD2. Subsequently, cells were incubated with PLA probes for 1 hour in a humidified chamber at 37°C. If the two oligo probes are in proximity, ligation occurs, and further amplification steps enable fluorescence detection of the reaction as bright red dots (566–594 nm). Cells were then counterstained with DAPI to visualize nuclei and imaged using Cytation10 widefield microscopy or leica stellaris confocal. Quantification of PLA dots were performed using Gen5 or ImageJ software.

#### Confocal immunofluorescence

Cells were fixed with 4% paraformaldehyde in PBS for 30 min at room temperature, treated with 0.1 M glycine for 10 min, and subsequently blocked/permeabilized with blocking buffer (PBS containing 1% BSA and 0.1% Triton X-100) for 20 min at room temperature. Primary and secondary antibodies were incubated for 1 h at room temperature or overnight in blocking buffer. Dilutions of antibodies used were as follows: HA (1:200) and DAPI (1:1000). Alexa fluor fluorescent dye conjugated secondary antibodies were used at 1:500 dilutions. For visualizing actin organizations cells were stained with 16 mM Phalloidin Alexa Fluor^TM^-594 for 30 min at room temperature, washed three times with PBS. ProLong Glass antifade reagent is used for mounting the coverslips on glass slides.

#### Immunofluorescence staining of colon tissue sections

Mouse colon Swiss rolls were fixed in zinc paraformaldehyde to prepare FFPE tissue blocks. FFPE tissue sections of 4 μm thickness underwent deparaffinization and rehydration. Heat-induced epitope retrieval was performed using Tris-EDTA buffer (pH 9.0) in a pressure cooker. Tissue sections were blocked using 2.5% goat serum, followed by incubation with primary antibody overnight in a humidified chamber at 4°C. Primary antibodies used for immunostaining were CD68 (mouse monoclonal, dilution 1:150, Abcam ab201340) and CD163 (rabbit polyclonal, dilution 1:150, Abcam ab182422). Goat anti-mouse IgG (H+L) AlexaFluor488 and Goat anti-rabbit (H+L) Alexa Fluor 594 were used as secondary antibodies (dilution 1:500) in combination with DAPI (dilution 1:1000) staining. Images were acquired on a Stellaris 5 Confocal Microscope (Leica microsystems) and images were analyzed using the software package QuPath (Version: 0.5.0) and further processed using Fiji/ImageJ software (NIH, Bethesda, USA).

#### CD68 and CD163 quantification

IF images were obtained in a random, genotype-blinded fashion and analyzed using QuPath (Version: 0.5.0). For each image, nuclei were detected based on DAPI and cell boundaries were drawn using a uniform 5um expansion with the QuPath built-in cell detection feature. An object classifier was created for the identification of CD163+, CD68+, and negative cell populations. Percent positivity was calculated by dividing the number of positive cells by the total number of cells in each image. Welch’s two sample unpaired t-test was performed to compute p values.

#### NFκB reporter assay

RAW 264.7 cells (50000 cells/well in 96-well plate) were transfected together with 50 ng NFκB reporter plasmid and 5 ng Renilla luciferase control plasmid. After 24h, cells were stimulated with L18-MPD (10 µg/ml) for 6 hr and NFκB activity was assessed using the Dual-luciferase Reporter Assay System using manufacturers protocol. For HeLa cells 10000 cells/well in 96-well plate were seeded and transfected with 25 ng NFκB reporter plasmid, 0.5 ng Renilla luciferase control plasmid and either 5 ng/well of NOD2-WT or its mutants. Cells were primed with or without L18-MPD (10 µg/ml) for 16 hr before stimulating with LPS100ng/ml for 6h and NFκB activity was assessed using a noncommercial dual luciferase enzyme assay as described (113).

#### RNA extraction and Quantitative PCR

Total RNA was isolated using TRIzol reagent (Life Technologies) and Quick-RNA MiniPrep Kit (Zymo Research, USA) as per manufacturer’s guidelines. RNA was converted into cDNA using the qScript™ cDNA SuperMix (Quantabio) and quantitative RT-PCR (qPCR) was carried out using PowerUp™ SYBR™ green master mix (Applied Biosystems, USA) with the StepOnePlus Quantitative platform (Life Technologies, USA). The cycle threshold (Ct) of target genes was normalized to 18S rRNA gene and the fold change in the mRNA expression was determined using the 2^-ΔΔCt method.

#### Gentamicin Protection Assay

Gentamicin protection assay was used to quantify viable intracellular bacteria as described previously (92). Approximately, 2 × 10^5^ TGPMs were seeded into 12-well culture dishes overnight before infection at an MOI of 10 for 1 h in antibiotic-free RPMI media containing 10% FBS in a 37 °C CO2 incubator. Cells were then incubated with gentamicin (200 μg/ml) for 90 min after PBS was to kill extracellular bacteria. After incubation time cells were washed with PBS and subsequently lysed in 1% Triton-X 100, lysates were serially diluted and plated on LB agar plates. Bacteria colonies (CFU) were counted after overnight (16 h) incubation at 37 °C. To test effect of MDP, cells were pre-treated overnight with MDP (10 µg/ml).

#### Cytokine Assays

Cytokines including TNFA, IL6, IL1B and IL10 were measured in cell supernatant using ELISA MAX Deluxe kits from Biolegend as per manufacturer’s protocol.

#### Statistics and reproducibility

All experimental values are presented as the means of replicate experiments ±SEM. Statistical analyses were performed using Prism 9 (GraphPad Software). Differences between the two groups were evaluated using Student’s t-test (parametric) or Mann-Whitney U-test (non-parametric). To compare more than three groups, one-way analysis of variance followed by Tukey’s post-hoc test was used. Differences at P < 0.05 were considered significant.

#### Study approval

All mice studies were approved by the University of California, San Diego Institutional Animal Care and Use Committee (IACUC) under protocol number S17223.

## Data availibility

The code related to the computational analyses used in this paper is available at: https://github.com/sinha7290/NOD2. Mass spectrometry proteomics data have been deposited in the ProteomeXchange Consortium via the PRIDE (114) partner repository under the dataset identifier PXD066180. All data supporting the findings of this study are included in the Supporting Data Values file, and complete, unedited blots are in the supplemental material.

## Author contributions

G.D.K, M.A and P. G. conceptualized the project. G.D.K., M.A., S.S, ME, EM, K.C-P, S-T.H, J.C, Y.M, C.R.E, M.M and V.C, were involved in data curation and formal analysis. D.T.V. carried out 16S microbiome analysis. S.S, with supervision from P.G, carried out all the transcriptomic and proteomic analyses. G.D.K conducted all animal studies. M.A conducted all biochemical studies. V.C assisted G.D.K. for ELISA. Y.M assisted G.D.K. for qPCR and image analyses. C.R.E and S-R.I assisted M.A in construct design, cloning and mutagenesis. G.D.K. and P.G. prepared Figures for data visualization, wrote the original draft, reviewed and edited the manuscript. P.G. supervised and acquired funding to support the study. All co-authors approved the final version of the manuscript.

## Acknowledgments

This work was supported by the National Institutes of Health (NIH) grant (R01-AI141630, UG3TR003355, UG3TR002968, and R01-AI55696), a pilot award from the Propel a Cure Foundation and the Leona M. and Harry B. Helmsley Charitable Trust (to P.G.). G.D.K. received support from the American Association of Immunologists Intersect Fellowship Program for Computational Scientists and Immunologists and the American Heart Association Career Development Award (24CDA1268506). M.S.A. was supported by an American Heart Association Predoctoral Fellowship (25PRE1357971). S.S. was supported by The AAI Intersect Fellowship Program. Additional support includes T32GM8806 (to D.V.). S.-R.I. received a postdoctoral fellowship from the NIH (3R01DK107585-02S1), and M.M. was supported by a UC San Diego Agilent Center of Excellence Postdoctoral Fellowship. The authors thank the UC San Diego Cellular and Molecular Medicine Electron Microscopy Core (RRID: SCR_022039) for access to equipment and technical assistance; the Core is partially supported by NIH grant S10-OD023527. Data in this manuscript were generated at the UC San Diego Institute of Genomic Medicine using an Illumina NovaSeq 6000, funded by NIH SIG grant S10-OD026929. The authors acknowledge instrumentation resources at the UC San Diego Agilent Center of Excellence in Cellular Intelligence. The content is solely the authors’ responsibility and does not necessarily represent the official views of the Helmsley Charitable Trust or the NIH.

