## Supplementary information for "Distinct Colitis-Associated Macrophages Drive NOD2-Dependent Bacterial Sensing and Gut Homeostasis"

**Affiliations:**
[1]Department of Cellular and Molecular Medicine, University of California San Diego, CA, USA.
[2]Department of Pathology, University of California San Diego, CA, USA.
[3]Department of Pediatrics, University of California San Diego, CA, USA.
[4]Department of Medicine, University of California San Diego, CA, USA.

[†] Equal contribution

**Conflict of interest statement:** Authors have declared that no conflict of interest exists.

**KEY WORDS**: GIV/Girdin, Guanine-nucleotide exchange modulators (GEMs), CCDC88A, Macrophage, NOD2, MDP, Microbes, Innate immunity

***Correspondence to**:

**Pradipta Ghosh, M.D.;** Professor, Departments of Medicine, and Cell and Molecular Medicine, University of California San Diego; 9500 Gilman Drive (MC 0651), George E. Palade Bldg, Rm 232, 239; La Jolla, CA 92093. Phone: 858-822-7633: Fax: 858-822-7636:

#### CATALOG OF SUPPLEMENTARY MATERIALS

1. *Supplementary Table (S1)*
2. *Supplementary Figures and Legends (S1-S4)*

#### SUPPLEMENTARY TABLES

##### Supplementary Table 1: NOD2 mutations and its clinical relevance in CD patients.

| Mutation | Percentage of CD Patients | % CD diagnosis | Mechanistic insights | Variation Caused | Phenotypic Consequence |
|---|---|---|---|---|---|
| **1007fs (Frameshift Stop codon mutation)** | 31% | 100% | Impaired palmitoylation (Lu et al. 2019) , and protein mislocalization<br><br>Forced PM-localization of protein *does not* restore function (Lécine et al. 2007)<br><br>MDP-binding site unaffected<br><br>Loss of binding to GIV (current work) | - Mutant is unable to recognize MDP to initiate NFkB activation<br>- Associated defective release of IL-10 from blood mononuclear cells (PBMC) after stimulation with the TLR2 ligands, PGN and Pam3Cys-KKKK, and LPS. | - The genotype relative risk (GRR) for developing CD in heterozygotes and homozygotes of this mutation alone is 3.29 ±0.64 and 34.66 ±12.87 respectively<br><br>- Homozygosity is strongly associated with gastroduodenal CD and younger age at diagnosis<br><br>- Homozygous patients demonstrate a much more severe disease phenotype than other patients with Crohn's disease and have an increased risk for ileal stenoses and surgical interventions |
| **R702W (Missense Substitution Mutation)** | 32% | 100% | Impaired palmitoylation (Lu et al. 2019) , and protein mislocalization<br><br>Forced PM-localization of protein restores function (Lécine et al. 2007)<br><br>MDP-binding site unaffected<br><br>Loss of binding to GIV (current work) | - Monocyte-derived dendritic cells (MoDCs) produced significantly higher levels of IL-12 on stimulation with whole bacteria<br><br>- MoDCs carrying the mutation displayed an increased basal level of IL-8 release, which, after a bacterial encounter, equilibrated to the levels similar to healthy controls. | - GRR for developing CD in heterozygotes and homozygotes of this mutation alone is 1.97±0.85 and roughly 3.05 respectively<br><br>- Positive independent association with structuring behavior and granuloma formation |
| **G908R (Missense Substitution Mutation)** | 18% | 80% | Normal palmitoylation (Lu et al. 2019) and protein mislocalization.<br><br>Forced PM-localization of protein restores function (Lécine et al. 2007)<br><br>One of the residues that form contact site for MDP and hence, MDP recognition could be directly impacted (Vijayrajratnam et al. 2017). GIV binding (current work). | -MoDCs produced significantly higher levels of IL-12 on stimulation with whole bacteria | - GRR for developing CD in heterozygotes and homozygotes of this mutation alone is 1.97±0.85 and 4.55±1.34 respectively |

### SUPPLEMENTARY FIGURES

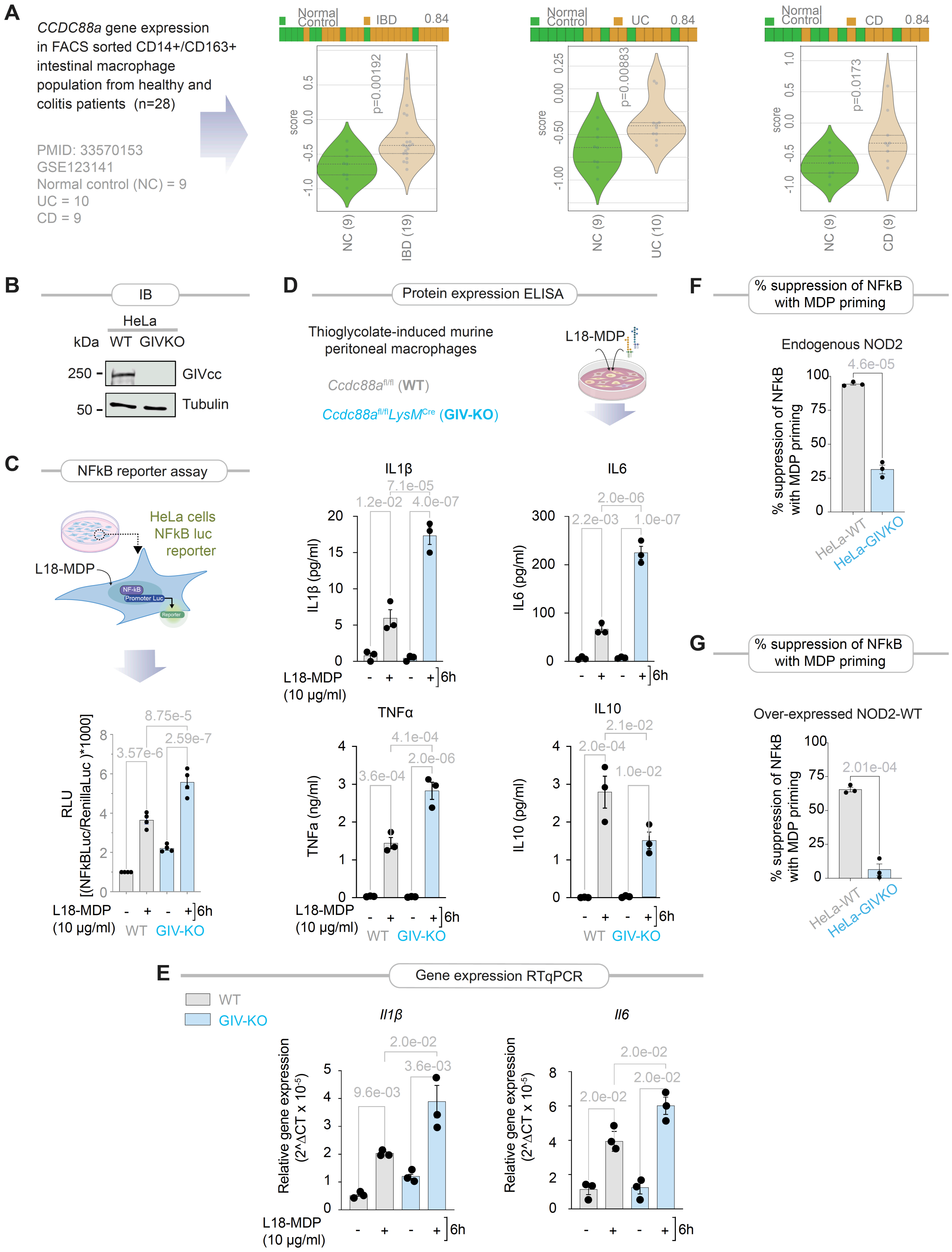


**Supplementary Figure 1: GIV is required for MDP-stimulated inflammatory resolution.** Related to **Figure 1, 2 and 5**.

**A.** Violin plots display *CCDC88A* expression in macrophages that were isolated from the lamina propria of colons from healthy and IBD (UC and CD) subjects. *p*-values are calculated using Welch's t-test. Bar plots on top denote sample classification accuracy using *CCDC88A* as single gene expression score and the numbers on top indicate ROC AUC.

**B.** Immunoblot (IB) of control (HeLa-WT) or GIV-KO (HeLa-GIV-KO) HeLa cells confirming depletion of GIV by >95%.

**C**. Schematics display the NFkB reporter assay in HeLa cells (*top*) and bar graphs display the fold change in NFkB activity (*bottom*).

**D.** Schematic (top) of study design to assess cytokines produced by peritoneal macrophage stimulated with L18-MDP (10 µg/ml) for 6 h. Bar graphs (bottom) display the concentrations of the indicated cytokines, as determined by ELISA.

**E.** Bar graphs display the gene expression in cell lysates in D, as determined by RT qPCR.

**F-G**. Bar graphs display the fold change in NFkB activity induced by LPS (100 ng/ml) for 6h, following 24-hour priming with L18-MDP (10 µg/ml) in HeLa cells. Panel F represents cells with endogenous NOD2, while Panel G represents cells with overexpressed NOD2.

*Statistics*: All results are displayed as mean ± SEM (n=3 biological replicates). Statistical significance was tested using two-way/one-way ANOVA followed by Tukey's test for multiple comparisons. *p*-value ≤ 0.05 is considered as significant.

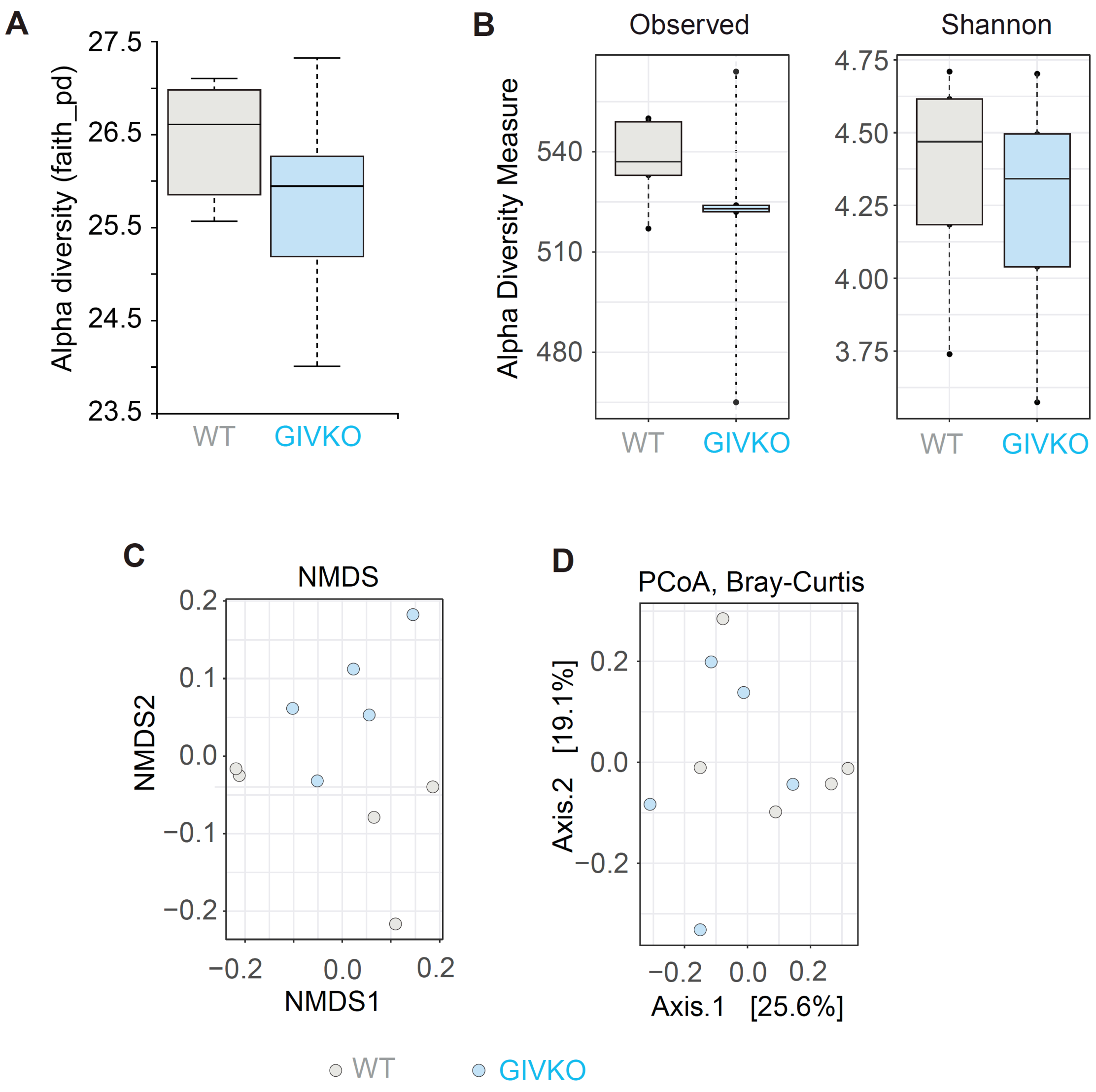


**Supplementary Figure 2: Fecal microbiome analysis confirms spontaneous dysbiosis in myeloid-specific (LysMCre) GIV-KO mice age 8-12 wk.** Related to **Figure 4A**.

**A-B**. Box plots display alpha diversity indices (Observed and Shannon), of fecal microbiome communities within GIV-KO and their littermate controls.

**C**. Non-Metric Multidimensional Scaling (NMDS) ordination of fecal microbiome communities within GIV-KO and their littermate controls

**D**. PCoA plot of beta diversity of fecal microbiome communities within GIV-KO and their littermate controls.

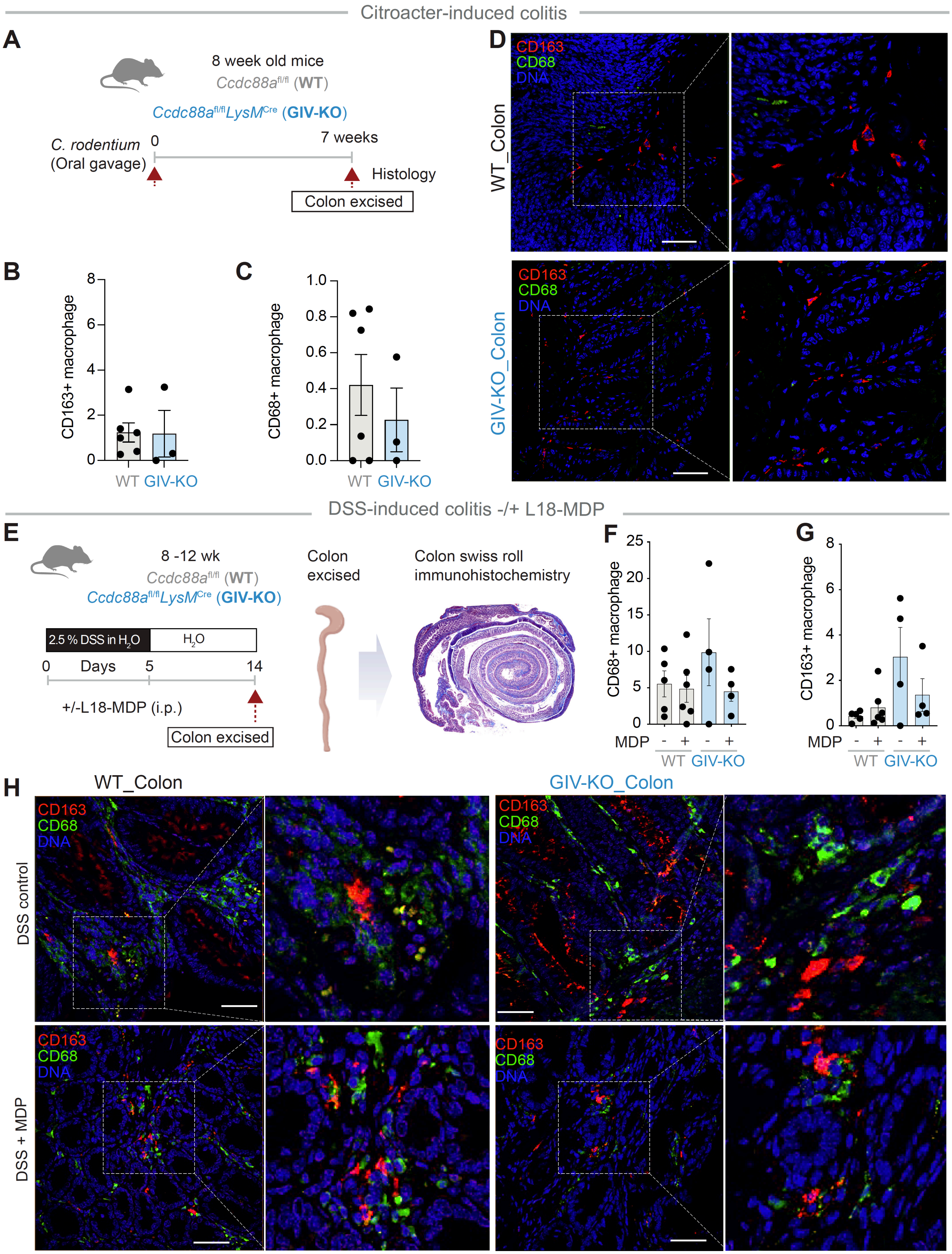


**Supplementary Figure 3: The Loss of GIV does not impede macrophage recruitment and expression of conventional markers for polarization.** Related to **Figures 4-5**.

**A-D.** Panels describing the experimental design (**A**) and macrophage population distribution in the colon (**B-D**) in an infectious colitis model of GIV-KO and control littermates induced using *Citrobacter rodentium* (initially

termed *Citrobacter freundii* biotype 4280 (1); strain name DBS100; 5 x $10^8$ CFU/200ul/mouse. GIV-KO, n=8; WT, n=6. Findings are representative of two independent repeats. *See Figure 4C-I for the details*.
**B-C.** Quantification of $CD163^+$ (B) and $CD68^+$ (C) macrophages in colonic sections. n=4-6. **D.** Representative immunofluorescence images showing CD163 (red), CD68 (green), and nuclei (blue) in colon tissues. Scale bars = 50 µm.
**E–H.** Schematic (E) displays the study design for DSS-induced colitis. GIV-KO, n=5; WT, n=5. Findings are representative of two independent repeats. *See Figure 5A-E for the details.*
**(F–G)** Quantification of $CD68^+$ (F) and $CD163^+$ (G) macrophages in colon tissues. n = 5–6 mice per group.
**(H)** Representative immunofluorescence images of colon sections from DSS-only and DSS+MDP groups showing CD163 (red), CD68 (green), and nuclei (blue). Scale bars = 50 µm.
All graphs represent mean ± SEM. Statistical comparisons were made using unpaired t-tests. $p$-value ≤ 0.05 is considered as significant.

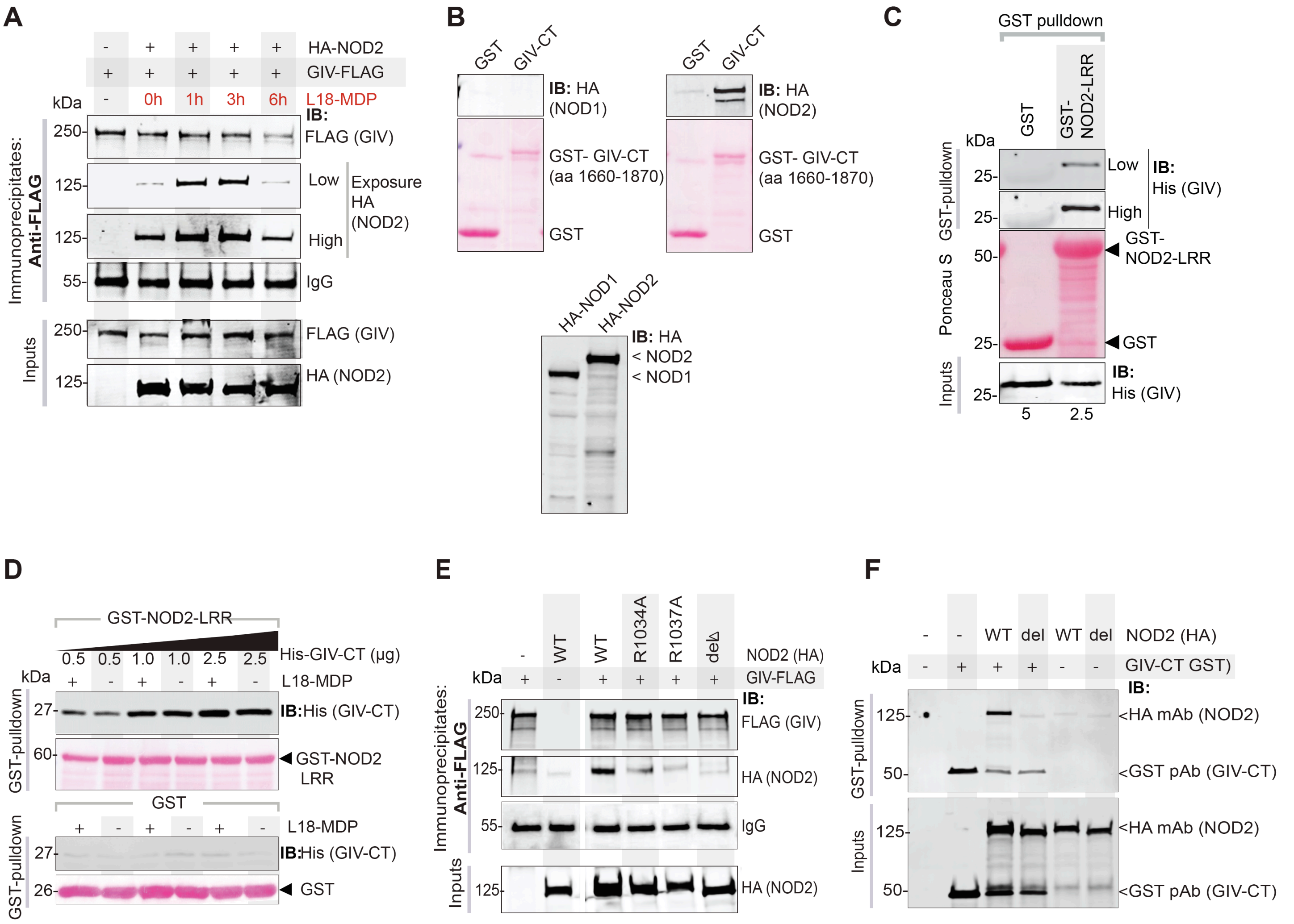


**Supplementary Figure 4: The NOD2(LRR)•GIV(C-term) interaction is a dynamic.** Related to **Figures 7-8**.

**A**. FLAG-tagged GIV was immunoprecipitated with anti-FLAG mAb from equal aliquots of lysates of HEK cells expressing GIV-FLAG and HA-NOD2, stimulated (+) or not (-) with L18-MDP (10 µg/ml) for indicated time points. Immunoprecipitated (IP; top) complexes and input (bottom) lysates were analyzed for NOD2 and GIV by immunoblotting (IB).

**B.** Lysates of HEK cells expressing HA-tagged NOD1/2 proteins were used in a GST pulldown assay with GST or GST-GIV-CT immobilized on Glutathione Sepharose beads. Bound NOD1/2 proteins were visualized by immunoblotting (IB).

**C.** Recombinant His-GIV-CT proteins were used in a GST pulldown assay with GST or GST-NOD2-LRR immobilized on Glutathione Sepharose beads. Bound GIV were visualized by immunoblotting (IB).

**D.** Recombinant His-GIV-CT proteins (0.5, 1, 2.5 µg) were used in a GST pulldown assay with GST or GST-NOD2-LRR immobilized on Glutathione Sepharose beads, in presence (+) or absence (-) of 10x molar excess of L18-MDP. If MDP and GIV share the same binding site on the LRR domain of NOD2, excess of MDP is expected to compete with GIV and reduce its binding to NOD2-LRR. Bound GIV was visualized by immunoblotting (IB).

**E**. FLAG-tagged GIV was immunoprecipitated with anti-FLAG mAb from equal aliquots of lysates of HEK cells expressing GIV-FLAG and either wild-type (WT) or mutant HA-NOD2 constructs. Immunoprecipitated (IP; top) complexes and input (bottom) lysates were analyzed for NOD2 by immunoblotting (IB).

**F**. GST-GIV-CT was pulled down using Glutathione Sepharose beads from equal aliquots of lysates of HEK lysates expressing the wild-type (WT) or del mutant HA-NOD2 construct either alone (last two lanes) or with GST-GIV-CT (aa 1660-1870; mammalian p-CEFL vector). Bound NOD2 proteins and similar expression of GIV-CT was assessed by immunoblotting (IB) using anti-HA (NOD2) and anti-GST (GIV-CT) antibodies.

**Uncategorized References**

1. Newman JV, Zabel BA, Jha SS, and Schauer DB. Citrobacter rodentium espB is necessary for signal transduction and for infection of laboratory mice. *Infect Immun.* 1999;67(11):6019-25.
